# Feasibility of enhancing carbon sequestration and stock capacity in temperate and boreal European forests via changes to management regimes

**DOI:** 10.1101/2022.02.10.479900

**Authors:** D. Dalmonech, G. Marano, J.S. Amthor, A. Cescatti, M. Lindner, C. Trotta, A. Collalti

**Affiliations:** Forest Modelling Lab., Institute for Agriculture and Forestry Systems in the Mediterranean, National Research Council of Italy (CNR-ISAFOM), Via Madonna Alta 128, 06128, Perugia, Italy; Forest Ecology, Institute of Terrestrial Ecosystems, Department of Environmental Systems Science, ETH Zurich, Zurich, Switzerland; Center for Ecosystem Science and Society, Northern Arizona University, Flagstaff, Arizona, 86001, USA; European Commission, Joint Research Centre (JRC), Ispra, Italy; European Forest Institute, Platz der Vereinten Nationen 7, 53113, Bonn, Germany; Department of Innovation in Biological, Agro-food and Forest Systems (DIBAF), University of Tuscia, 01100 Viterbo, Italy; Department of Agriculture and Forest Sciences, University of Tuscia, 01100 Viterbo, Italy

**Keywords:** adaptive forest management, modeling, virtual forests, climate change, carbon sequestration

## Abstract

Forest management practices might act as nature-based methods to remove CO_2_ from the atmosphere and slow anthropogenic climate change and thus support an EU forest-based climate change mitigation strategy. However, the extent to which diversified management actions could lead to quantitatively important changes in carbon sequestration and stocking capacity at the tree level remains to be thoroughly assessed. To that end, we used a state-of-the-science bio-geochemically based forest growth model to simulate effects of multiple forest management scenarios on net primary productivity (NPP) and potential carbon woody stocks (pCWS) under twenty scenarios of climate change in a suite of observed and virtual forest stands in temperate and boreal European forests. Previous modelling experiments indicated that the capacity of forests to assimilate and store atmospheric CO_2_ in woody biomass is already being attained under business-as-usual forest management practices across a range of climate change scenarios. Nevertheless, we find that on the long-term, with increasing atmospheric CO_2_ concentration and warming, managed forests show both higher productivity capacity and a larger potential pool size of stored carbon than unmanaged forests as long as thinning and tree harvesting are of moderate intensity.

## 1 Introduction

Forest ecosystems have the capacity to slow anthropogenic climate change by reducing the rate of atmospheric carbon dioxide (CO_2_) increase through increased photosynthetic CO_2_ assimilation, tree growth, and carbon storage in plants and soil, including growth of wood that is harvested and converted into products such as paper and timber (Pugh et al., 2019; Friedlingstein et al., 2020). Reducing net CO_2_ emissions is the central priority for achieving climatic stability in Europe by the target date of 2050 (European Green Deal). A key research question is whether current or future forest management practices can provide a concrete, cost-effective toolset for enhanced carbon storage at the ecosystem and landscape scales (Kauppi et al., 2001; Pussinen et al., 2002; Nolè et al., 2015; Tong et al., 2020) as well as greater wood harvesting for material substitution purposes (Leskinen et al., 2018; Howard et al., 2021).

European (EU) forests have been shaped by centuries of human activities that affect their present potential for carbon storage (Nabuurs et al., 2008). In the EU today, *circa* 165 Mha of forests are managed in ways that drive a net uptake of about 286 Mt CO_2_ year^−1^ (Grassi et al., 2017, 2021). Past management strategies were designed to maximize yield of wood products (Leslie, 1966) rather than total biomass production or carbon storage (Tahvonen, 2016). Recent EU-level policies are directed toward sustainable and climate-resilient forests (MCPFE 1990-2003; Forest Europe 2010; Forest Europe 2015; EU Forest Strategy 2015; State of Mediterranean Forest 2018, 2020; New EU-Forest Strategy; 2021; EU Adaptation Strategy 2021) to secure ecosystem services (including carbon sequestration for climate change mitigation) under changing climatic conditions (Churkina et al., 2020; Favero et al., 2020). However, substantial uncertainty remains about the effectiveness of current management practices to maintain the current carbon sink under ongoing climate changes, and their ability to enhance carbon storage in the future, because classical silvicultural schemes were developed in the context of past environmental conditions (Bellassen et al., 2011). In addition, in recent decades European forests achieved increased productivity that exceeded harvesting rates (Ciais et al., 2008; State of Europe’s forests 2020). These productivity increases resulted from the combinations of several factors: changes to forest age distribution, climate change (e.g. lengthening of the growing season; Peano et al., 2019), increased atmospheric CO_2_ concentration (stimulating photosynthesis through ‘CO_2_-fertilization’; Chen et al., 2022), increased nitrogen deposition (stimulating growth through nitrogen fertilization; Etzold et al., 2020; Chen et al., 2022) and forest management (Bellassen et al., 2011; Piao et al., 2020; Walker et al., 2021). But these positive trends may not continue at rapid rates in the future and could be offset by increased disturbance frequency and severity (Seidl et al., 2017; Senf and Seidl, 2021a), raising concerns that recent harvesting rates may be approaching, or even exceeding, net tree growth rates in temperate and boreal European forests (Nabuurs et al., 2013; Ceccherini et al., 2020; Schulze et al., 2020; State of Europe’s forests 2020). Moreover, recent positive trends in gross primary productivity (GPP; photosynthetic assimilation of atmospheric CO_2_) and plant growth in the northern hemisphere might not be sustained in the future if the CO_2_ fertilization effect is not persistent (Körner, 2005; Walker et al., 2021; Wang et al., 2021) or if they are counteracted on by increased heat stress and drought (Yuan et al., 2019; Grossiord et al., 2020; Wang et al., 2020), disturbance-related tree mortality (McDowell et al., 2020; Senf and Seidl, 2021b; Gampe et al., 2021; Hlasny et al., 2021), or by tree-age-related effects on net primary productivity (NPP; the balance of photosynthetic CO_2_ assimilation and plant respiratory CO_2_ release) (Ryan et al., 2006; Zaehle et al., 2006; Luyssaert et al., 2007; Tang et al., 2014; Pugh et al., 2019). Hence, a reduction in the forest net carbon sink capacity and its transition to a CO_2_ source could be approaching (Duffy et al., 2015; Peñuelas et al., 2017; Wang et al., 2020). This might reduce both the potential for forests to sequester additional carbon and sustainably produce wood products.

Forest carbon cycle and productivity models are used to investigate and project the effects of climate change and management scenarios on forest productivity across a range of spatial and temporal scales to support science-driven policymaking (Temperli et al., 2012; Vacchiano et al., 2012; Reyer et al., 2017; Maréchaux et al., 2021). This includes design of alternative management strategies to support climate change mitigation policies (Maréchaux et al., 2021; De Marco et al., 2022).

This study aimed to inform the debate about the role of forest management practices in sustaining high forest productivity under future climate conditions. To that end, a validated process-based modeling approach was designed to quantify controls on CO_2_ uptake and C storage in a composite matrix of managed forests, taking into account how combinations of climate change and management scenarios may affect those controls. Specifically, we questioned whether, relative to business-as-usual (BAU) management scenarios, alternative forest management practices could maximize NPP while at the same time maintaining and/or increasing potential Carbon Woody Stocks (pCWS; i.e., when no decay of harvested wood products occurs) in response to a range of climate change scenarios.

## 2. Materials and Methods

### 2.1 3D-CMCC-FEM Model

#### 2.1.2 Model description

The 3D-CMCC-FEM v.5.5 (Three Dimensional - Coupled Model Carbon Cycle - Forest Ecosystem Model; Collalti et al., 2014, 2016; Marconi et al., 2017; Engel et al., 2021; Mahnken et al., 2022) simulates daily GPP through the Farquhar-von Caemmerer-Berr photosynthesis model (Farquhar et al., 1980), modified for sun and shaded leaves (de Pury and Farquhar, 1997) and acclimation to temperature history (Kattge and Knorr, 2007). Plant respiration (R_a_) is simulated explicitly and partitioned into growth (R_g_) and maintenance (R_m_) fractions as in the growth-and-maintenance-respiration paradigm (Amthor, 2000; McCree, 1970; Thornley, 2000). R_m_ is computed, for each functional-structural tree C pool (i.e. live wood, leaves and fine roots), using a temperature-acclimated Q_10_ relationship (for details on thermal acclimation see Tjoelker et al., 2001; Atkin and Tjoelker, 2003; Smith and Dukes, 2012; Collalti et al., 2018) and a N-based maintenance respiration (m_R_) rate for living tissues of 0.218 g C g N^−1^ day^−1^ (Ryan et al., 1991; Amthor and Baldocchi, 2001; Oleson et al., 2013; Drake et al., 2016Collalti et al., 2016, 2020a). R_g_ is considered a fixed fraction (i.e. 30%) of the remaining C once tissue R_m_ is accounted for and removed from GPP. The sum of daily R_g_ (if any) and R_m_ gives R_a_. Daily NPP is then GPP less R_a_. Allocation of NPP among tree C pools is performed daily, with preference to a non-structural carbon pool (NSC, i.e. storage in starch and sugars), which is used directly to fuel R_m_, up to a minimum NSC threshold level. The minimum NSC-threshold level is a fraction (a model parameter) of the live wood C- content (Collalti et al., 2020a). Once (and if) the minimum NSC threshold is reached, C is allocated preferentially for biomass growth for the different tree structural C-pools depending on the phenological phase as formerly described in Collalti et al. (2016). The only phenological phase during which NSC has no priority in allocation is during bud break (D’Andrea et al., 2020, 2021), when recent GPP is completely allocated for growth of leaves up to a maximum annual leaf area index (LAI, m^2^ m^−2^), which is computed at the beginning of each year of simulation through the pipe-model (Shinozaki et al., 1964; Mäkelä, 1997), and growth of fine roots. This NSC allocation scheme reflects a quasi-active role of NSC, with NSC usually having priority over growth of new structural tissues, as described by Sala (2011), Merganičová et al. (2019) and Collalti et al. (2020a). This implies that any asynchrony between C-demand (i.e. respiration and growth) and C-supply (photosynthesis) is buffered by changes to the NSC pool. When NSC approaches zero and cannot be refilled by GPP, carbon starvation occurs, and tree death is simulated. This overall C-allocation scheme in the 3D-CMCC-FEM model follows the functional balance theory of allocation, similarly to other models (Merganičová et al., 2019). Age-related mortality, carbon starvation, and a background mortality (i.e. the as-yet unexplained mortality), represent the different types of mortality simulated by the model; the last one is turned off when forest management is applied. An in-depth description of the model’s underlying characteristics, responses to climate change, and sensitivity to model parameter values, as well as model limitations, is in Collalti et al. (2019, 2020a, and references therein).

#### 2.2.2 Forest management routine

Historically, the majority of European forests were actively managed and the share of coniferous species, particularly Scots pine and Norway spruce, was favored due to their high growth increment rates (Naudts et al., 2016). Management techniques (thinning and clear-cutting) aimed at maximizing productivity and reducing losses often resulted in even-aged, mono-specific forest stands (Campioli et al., 2015, and references therein; State of Europe’s forests 2020; Figure S1 in Supplementary Material). In such stands, carbon pools and fluxes strongly depend on rotation lengths (tree age-class distribution), thinning interval, and thinning intensity (Kaipainen et al., 2004; Nabuurs et al., 2008).

In this study we varied these three key management factors: thinning intensity, thinning interval and rotation age (following Reyer et al., 2020). Thinning intensity is represented in the model by the percentage of stand basal area to be removed based on the total stand basal area. Thinning interval is the number of years between two consecutive operations. Rotation age is the stand age at which a full harvest occurs, after which the stand is replanted with saplings of the species adopted in the Inter-Sectoral Impact Model Intercomparison Project (ISIMPI, https://www.isimip.org, Warszawski et al., 2014) protocol. The model benchmark was the BAU forest management scheme for the most common European species as described in Reyer et al. (2020) and Mahnken et al. (2022) and applied in three contrasting forest stands as in Collalti et al. (2018).

### 2.2. Sites, data and experimental design

The model was parameterized for, and simulated C fluxes and tree growth in three even-aged, long-monitored, managed European forest sites which are part of the Fluxnet network (Pastorello et al., 2020), the ISIMIP initiative and the PROFOUND database (Reyer et al., 2020; Mahnken et al. 2022): (1) the temperate European beech (*Fagus sylvatica* L.) forest of Sorø, Denmark; (2) the Norway spruce (*Picea abies* (L.) H. Karst) stand of Bílý Křìž in Czech Republic, and (3) the boreal Scots pine (*Pinus sylvestris* L.) forest of Hyytiälä, Finland (Table 1). These sites were selected because they represent the dominant forest types in Europe covering the temperate and boreal climatic zones (Figure S1),which contain more than 50% of European forest biomass (Avitabile et al., 2020), and their management best corresponds to ‘the intensive even-aged forestry’ as defined by Duncker et al. (2012).

**Table 1.** Site description for model initialization data (corresponding to the year 1997 for Sorø and Hyytiälä and 2000 for Bílý Kříž) to the real stands’ characteristics, and management variables used in simulations (see also Collalti et al., 2018; Reyer et al., 2020). Values in brackets represent bounds of variability (the maximum and the minimum absolute values) adopted for alternative management simulations. Re-planting information for the sites in the simulation experiments, according to ISIMIP protocol as in Reyer et al. (2020). The real stands refer to the monitoring sites in Sorø (*F. sylvatica*, Denmark), Bílý Kříž (*P. abies*, Czech Republic) and Hyytiälä (*P. sylvestris*, Finland).

| Species | DBH<br>(cm) | Age<br>(years) | Tree<br>height<br>(m) | Density<br>(trees ha<sup>-1</sup>) | Thinning<br>intensity<br>(% basal area) | Thinning<br>interval<br>(years) | Rotation age<br>(years) | Replanting<br>Species | Density<br>(trees ha<sup>-1</sup>) | Age<br>(years) | Tree<br>height<br>(m) |
| --- | --- | --- | --- | --- | --- | --- | --- | --- | --- | --- | --- |
| *Fagus*<br>*sylvatica* | 25 | 80 | 25 | 400 | 30 (20-40) | 15 (5-25) | 140 (120-160) | *Fagus*<br>*sylvatica* | 6000 | 4 | 1.3 |
| *Pinus*<br>*sylvestris* | 10.3 | 36 | 10 | 1800 | 20 (10-30) | 15 (5-25) | 140 (120-160) | *Pinus*<br>*sylvestris* | 2250 | 2 | 1.3 |
| *Picea*<br>*abies* | 7.1 | 16 | 5.6 | 2408 | 30 (20-40) | 15 (5-25) | 120 (100-140) | *Picea*<br>*abies* | 4500 | 4 | 1.3 |

Daily weather values input to the model for each site were derived from the climate scenario data from the ISIMIP Fast Track initiative based on the Climate Model Intercomparison Project 5 (CMIP5) in which five Earth System Models (ESMs; i.e.: HadGEM2-ES, IPSL- CM5A-LR, MIROC-ESM-CHEM, GFDL-ESM2M, and NorESM1-M) were driven by four Representative Concentration Pathways (RCPs) of atmospheric greenhouse gas concentration trajectories, namely RCP 2.6, RCP 4.5, RCP 6.0, and RCP 8.5 (Moss et al., 2010; van Vuuren et al., 2011). Daily meteorological forcing-data for each site used by 3D-CMCC-FEM were available as bias-corrected/downscaled variables (air temperature, precipitation, solar irradiance) and as non-corrected variables (relative humidity) according to Hempel et al. (2013). The RCP atmospheric CO_2_ concentration values for the period 2016 to 2500, based on Meinshausen et al. (2011) as described in Reyer et al. (2020), were used to drive the biogeochemical photosynthesis model with values varying at the end of this century from 421.4 μmol mol^−1^ (RCP 2.6) to 926.6 μmol mol^−1^ (RCP8.5).

#### 2.2.1. Virtual stands

Given that the European forested area is composed of a mix of differing-aged stands, and since forest C cycle processes may respond differently to climate factors at different ages (e.g. Collalti and Prentice, 2019; Collalti et al., 2020b; Huber et al., 2018, 2020; Migliavacca et al., 2021), we developed a Composite Forest Matrix (CFM) consisting of a mixture of stands of different age, structure and associated biomass. Starting from the real stands, we generated a prescribed number of virtual stands in order to obtain representative model outputs of a larger set of different age-classes (with their associated forest attributes) to cover an entire rotation period (∼140 years, depending on species), similar to the approach in in Bohn and Huth (2017). Because 3D-CMCC-FEM, in accordance with experimental evidence, is sensitive to the stand structure (see Collalti et al., 2019, 2020a), this procedure allows a robust assessment of effects of management practices across the full range of applicable stand ages . A CFM framework was then created by running at each site the model from 1997 to 2199 (to cover the entire rotation length for each species) under a contemporary climate scenario (no climate change), consisting of de-trended and repeated cycles of 1996-2006 historical weather, with fixed atmospheric CO_2_ concentration of 368.8 μmol mol^−1^ and BAU management practices. From each of these simulations data needed to reinitialize the model at every *rotation length*/10 were extracted (Figure 1). Thus, in total, ten virtual stands representing different age classes of the composite matrix were selected and included into the CFM. The management scenario analysis was carried out on this set of forest virtual stands.

**Figure 1.**
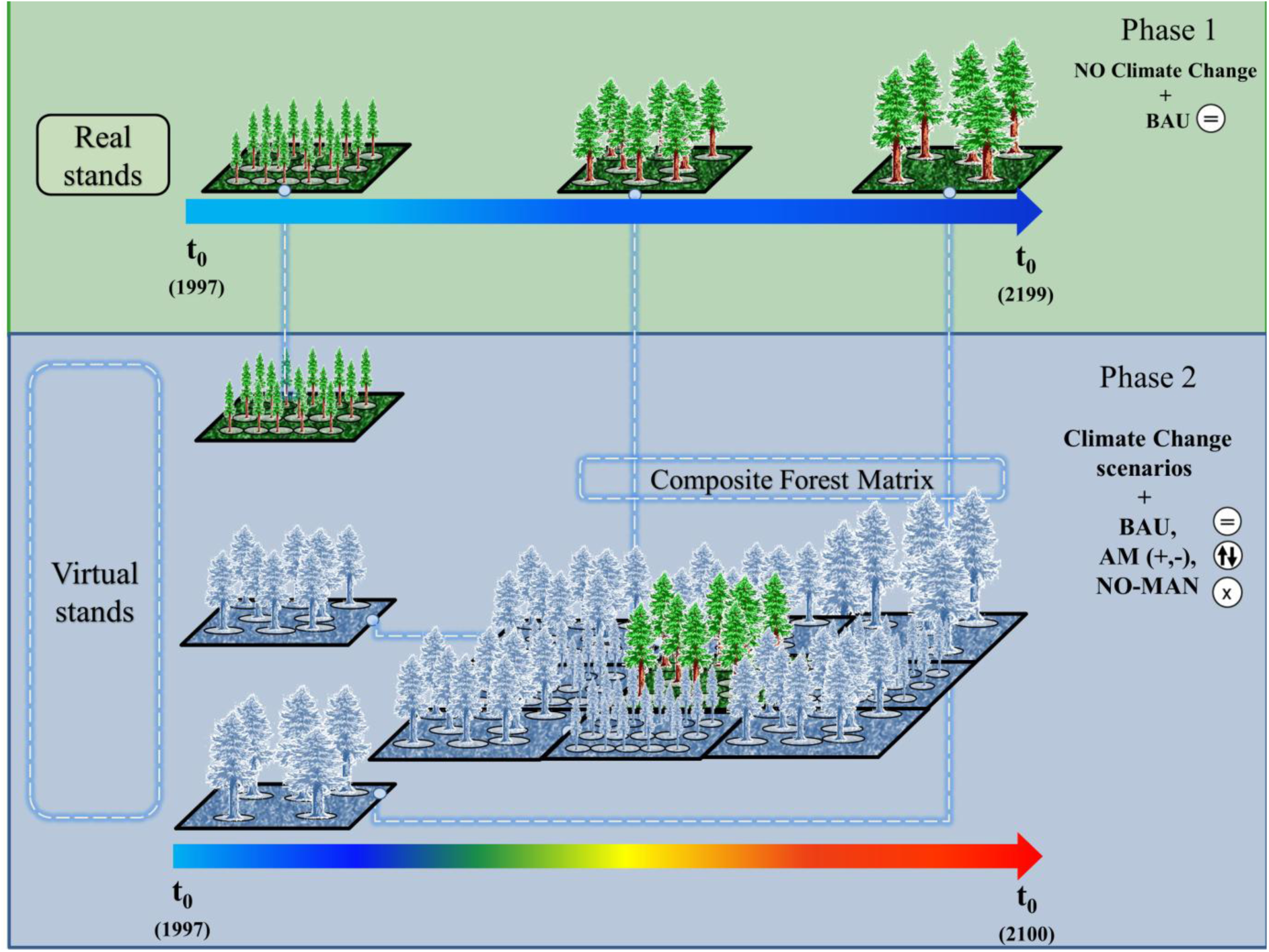
Conceptual scheme of the virtual stands creation: in Phase 1 the model is initialized with data from the actual forest stands and then simulations are carried out for 202 years of contemporary (1996-2006) weather and atmospheric CO_2_ concentration. In Phase 2, multiple stands are drawn from the simulations in Phase 1 and used to build the Composite Forest Matrix (CFM) composed of representative forest stands. The climate change (RCPs) and management scenarios (BAU, Alternative Managements, No-Management) simulations are then applied to the CFM.

#### 2.2.2. Alternative management (AM) schemes

For each real and virtual stand at each of the three sites we applied 28 management scenarios: the BAU and ‘no management’ (NO-MAN: stands allowed to grow without thinning or harvesting) schemes plus 26 alternative management schemes. These alternative forest management scenarios represent all the possible combinations of two thinning *intensities*, two thinning *intervals*, and two *rotation* durations than differ from those in the BAU scenario. The schemes were grouped (Tables 1 and Table S1) into combinations of: (1) ‘*more intensive*’ (‘AM+’), where at least one out of the three management variables reflect an intensified management case relative to BAU (e.g. higher thinning intensity and/or shortened interval and/or shortened rotation length than BAU), and the other one or two (or no) variables are kept as in BAU; (2) ‘*less intensive*’ (‘AM–’), where at least one variable reflects lower thinning intensity and/or prolonged interval and/or prolonged rotation length, compared to the BAU case; and (3) ’*mixed schemes*’ (‘MIX’), where at least one management variable was more intensive and at least one management variable was less intensive than the BAU scheme.

#### 2.2.3 Model runs and evaluation

The starting year for all simulations was 1997, consistently with the availability of measured stand carbon flux data used for the model initialization and evaluation. After creation of the virtual stands, based on de-trended historical weather time series for period 1996-2006, climate change simulations were conducted for the period 2006 to 2100. By considering all combinations of management and climate change scenarios from the different ESMs climate forcings, 6,160 simulations were performed at each site (i.e., 5 ESMs * 4 RCPs * 11 stands * 28 management schemes) for a grand total of 18,480 model runs.

To gauge model sensitivity to the study variables we organized the analysis according to a factorial design (Mason et al., 2003; Collalti et al., 2018) across a matrix of Stand, ESM and RCP, generating seven possible combinations from each factor. This was used to identify the most influential factor the modelled carbon cycle variables GPP, R_a_, NPP, carbon-use efficiency (CUE = NPP/GPP), net woody productivity (NPP_woody_), and potential Carbon Woody Stocks. The model was evaluated using 1997-2005 annual GPP and NPP_woody_ data for Sorø and Hyytiälä and 2000-2005 for Bílý Křìž by comparing simulated GPP against eddy covariance estimates (Pastorello et al., 2020), and compared modeled wood growth against measured NPP_woody_ (Principal Investigator’s site for Hyytiälä and Bílý Křìž under *personal communication*, and Wu et al., 2013 for the site of Sorø) and stem diameter at breast height (DBH; Reyer et al., 2020; Mahnken et al., 2022). Subsequent years were excluded from the model evaluation since the scenario period in the ESMs started in 2006, and hence was driven by different atmospheric CO_2_ concentration trajectories after 2006.

### 2.3 Effects of climate change and management on carbon fluxes and biomass

As carbon fluxes do not scale linearly to stocks (Schulze et al., 2020) data analyses focused separately on the variables NPP and pCWS, the sum of standing and previously harvested woody stocks. Net primary productivity can be a good proxy for forest carbon sink processes (Sha et al., 2022), and the net biomass input to forest ecosystems (Trotsiuk et al., 2020), with decomposition (decay, or heterotrophic respiration) processes representing the active carbon source process. Net primary production is a dynamic balance between photosynthesis and plant respiration, which respond separately and/or in combination to a range of climatic factors and, in managed forests, to management practices (Collalti et al., 2020a; 2020b). These age-dependent responses are generally not prone to *in situ* quantification over long periods, especially for climate change issues, hence raising the need for process-based modeling. Harvested wood products are considered here without decay; hence we aim at evaluating only the potential maximum attainable total woody standing stocks under a wide spectrum of possible management schemes without any consideration of the turn-over of harvested wood products. Data were averaged over the emission scenario simulation period 2006-2099 and aggregated over virtual and real stands and over ESMs but distinguished between RCPs. In spite of site-specific differences in magnitude of response to the different management schemes and climate (see Supporting Material) the main emerging pattern with increasing intensity of management intervention was of similar magnitude. For this reason, data were also aggregated over sites. Therefore, alternative management practice results are presented as aggregated according to the groups AM+ and AM– to highlight patterns of directional changes when moving from the current management schemes toward a more intensive or a less intensive scenario.

## 3. Results

### 3.1 Model evaluation

The 3D-CMCC-FEM model was evaluated at the three sites separately and at different temporal scales with robust data for both the carbon fluxes GPP and NPP_woody_, and the structural variable average stand DBH (Figures S2-S4 in Supplementary Material). Simulations forced with both observed local daily weather and with an ensemble of outputs from climate models for the contemporary period compare well with the eddy-covariance based estimated daily GPP values at the sites of Sorø (Root Mean Square Error, RMSE = 2.15 g C m^−2^ day^−1^ with local climate, and RMSE = 2.98 g C m^−2^ day^−1^ with the ensemble across ESMs forcing; r > 0.86), Hyytiälä (RMSE = 1.48 g C m^−2^ day^−1^ with local climate, and RMSE = 1.91 g C m^−2^ day^−1^ with the ensemble across ESMs forcing; r > 0.78) and Bílý Křìž (RMSE = 2.07 g C m^−2^ day^−1^ with local climate, and RMSE = 2.69 g C m^−2^ day^−1^ with the ensemble across ESMs forcing; r > 0.67) (see Supplementary Material Table S2-S3).

Modelled annual GPP was also consistent with site measurements at Sorø (1665 ± 171 g C m^−2^ year^−1^ and 1585 ± 190 g C m^−2^ year^−1^ modelled from observed and modelled climate vs. 1731 ± 184 g C m^−2^ year^−1^ measured; here and elsewhere, ± denotes one standard deviation)), Hyytiälä (894 ± 57 g C m^−2^ year^−1^ and 871 ± 52.6 g C m^−2^ year^−1^ modelled from observed and modelled climate vs. 1028 ± 50 g C m^−2^ year^−1^ measured), and Bílý Křìž (893 g C ± 252 g C m^−2^ year^−1^ and 893 ± 222 g C m^−2^ year^−1^ modelled from observed and modelled climate vs. 1024 ± 354 g C m^−2^ year^−1^ measured). Similarly, modelled values were similar to measured values of tree woody pools (i.e. NPP_woody_) at Sorø (351 ± 61 g C m^−2^ year^−1^ and 275 ± 63 g C m^−2^ year^−1^ modelled from observed and modelled climate vs. 346 ± 36 g C m^−2^ year^−1^ measured) and Bílý Křìž (442 ± 79 g C m^−2^ year^−1^ and 405 ± 36 g C m^−2^ year^−1^ modelled from observed and modelled climate vs. 380 ± 38 g C m^−2^ year^−1^ measured). The NPP_woody_ simulated at Hyytiälä was not quite as close to available data (317 ± 21 g C m^−2^ year^−1^ and 290 ± 24 g C m^−2^ year^−1^ modelled from observed and modelled climate vs. 228 ± 23 g C m^−2^ year^−1^ measured). Mean DBH increase, which was only qualitatively compared, is slightly underestimated at the site of Bílý Křìž (Figure S3). Comparisons of literature data for NPP, R_a_ and CUE with modelled values are in Table S3. Notably, results generated by3D-CMCC- FEM forced with the EMSs’ climates are close to those generated when forced with observed climate data in the evaluation period.

### 3.2 Less intensive management vs. BAU

Simulated average NPP in the less intensive management scenario group (i.e. AM–) is close to the reference BAU values, ranging from 495 g C m^−2^ year^−1^ (–1.2%; here and elsewhere, percentages refer to changes compared to BAU) to 525 g C m^−2^ year^−1^ compared to the range from 502 g C m^−2^ year^−1^ to 542 g C m^−2^ year^−1^ for RCP 2.6 and 8.5 under BAU, increasing slightly with warming and increased atmospheric CO_2_ concentration in the early years of the study period. Changes were larger toward the end of the century and without significant differences across RCPs (Figure 2 and Figure S4, Table 2). Simulated pCWS values increase steadily along the simulation time for all the alternative management scenarios, with time-averaged values in the range 180-199 t C ha^−1^ compared to 193-199 t C ha^−1^ for RCP 2.6 and 8.5 under BAU, respectively (Figure 3 and Figure S4, Table 2). Relative to BAU, the ratio pCWS/NPP decreased with less intensive management and with no management (Table 4; results for each of the RCPs scenarios and all the alternative management options combined are reported separately across the sites in the Supplementary Material Table S4).

**Figure 2.**
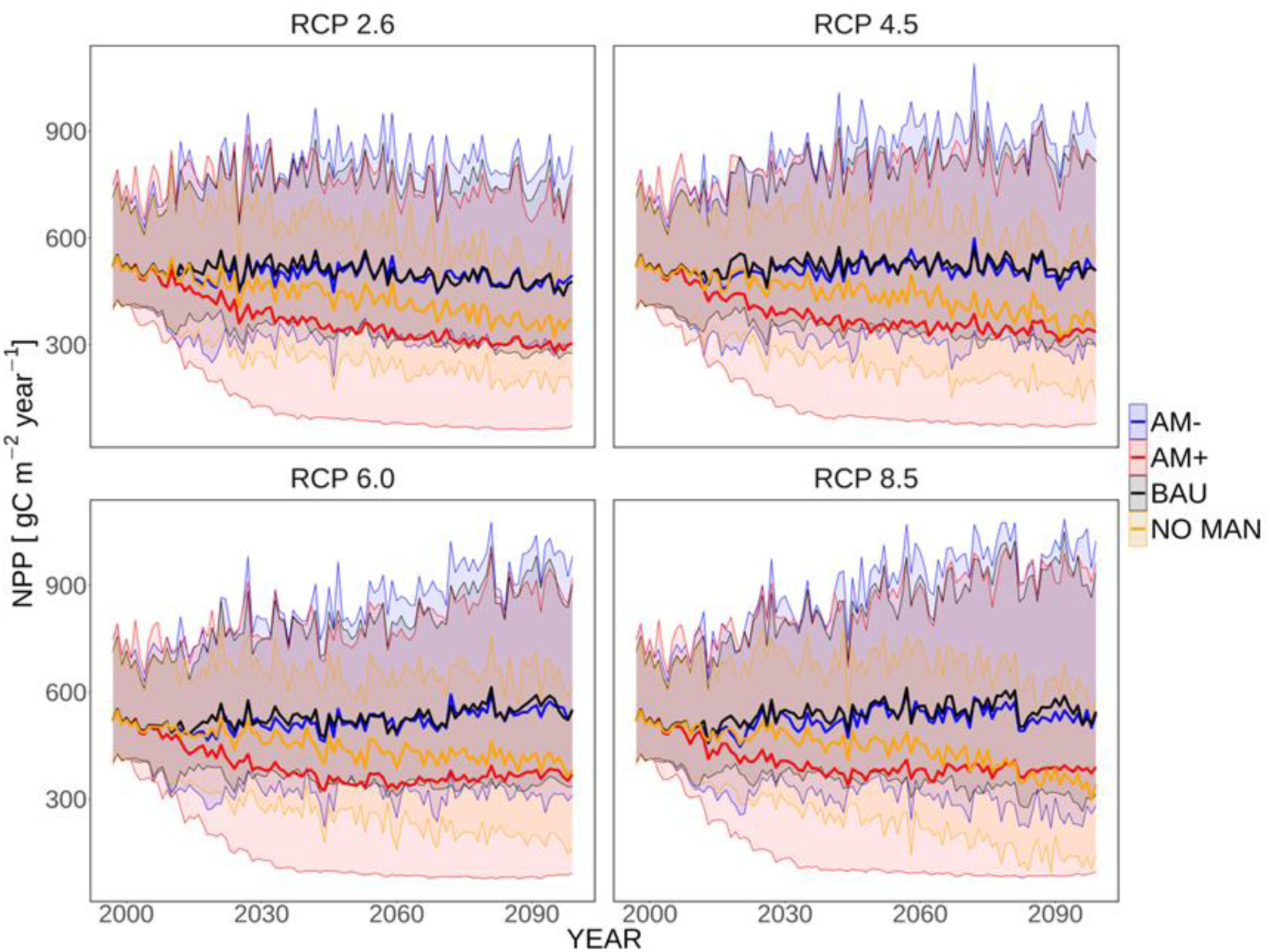
NPP (Net primary productivity, g C m^−2^ year^−1^) simulations under different management scenarios (AM+, BAU, AM–) and the NO-MAN scenario for each of the four atmospheric CO_2_ concentration pathways (RCPs). NPP, solid line, is averaged across the representative forests, different ESMs and aggregated according to the management regime. Shaded areas represent the maximum and minimum values (5^th^ and 95^th^ percentiles) across the representative forests, different ESMs and aggregated according to the management regime.

**Figure 3.**
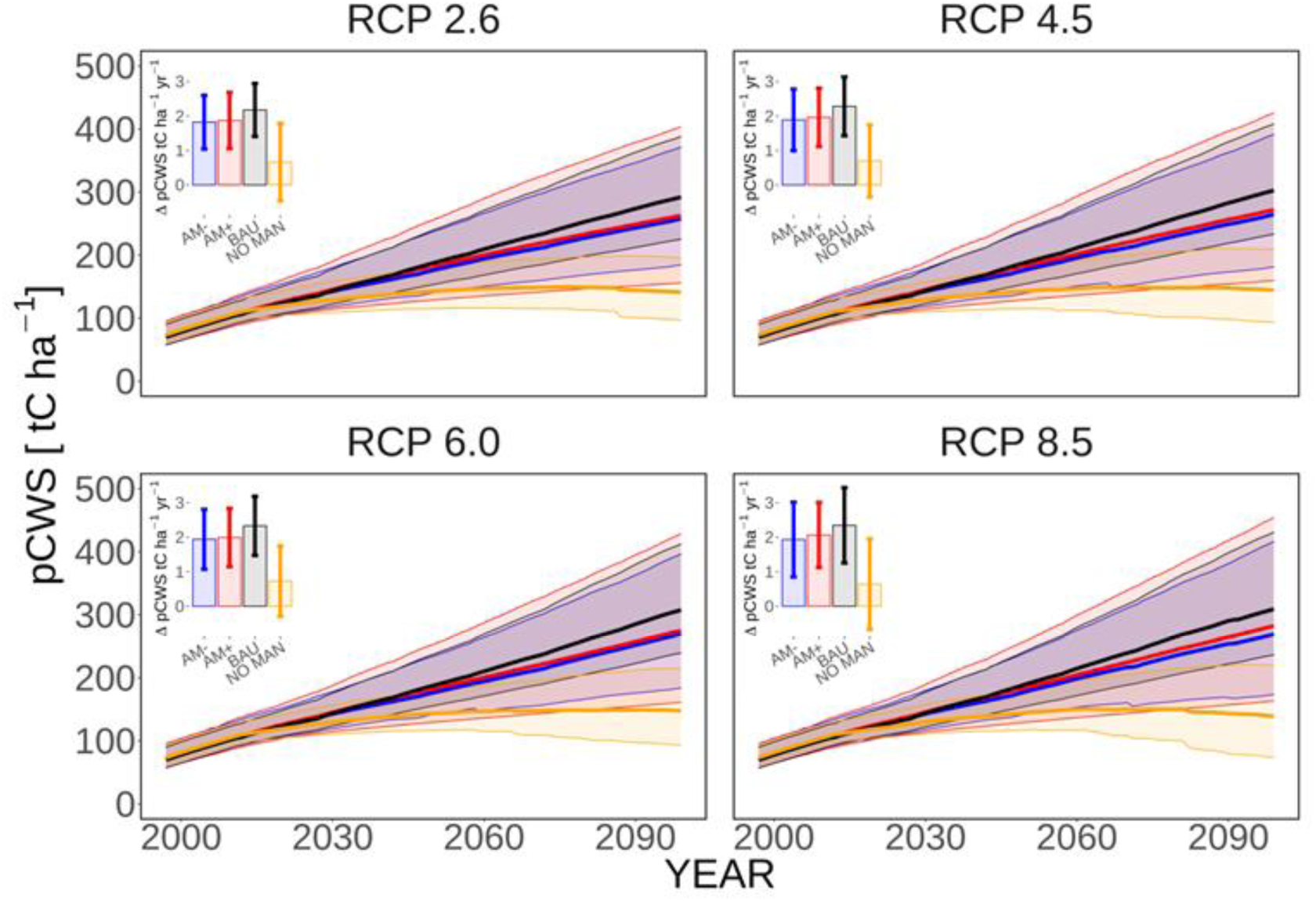
pCWS (potential Carbon Woody Stock = standing and potential harvested woody biomass; t C ha^−1^) simulations under different management scenarios (AM+, BAU, AM–) and the NO-MAN scenario divided by different emission scenario RCPs. pCWS, solid line, is averaged across the representative forests, different ESMs and aggregated according to the management regime. Shaded areas represent the maximum and minimum values (5^th^ and 95^th^ percentiles) across the representative forests, different ESMs and aggregated according to the management regime. Carbon sequestration rates (as annual increase of CWS, t C ha^−1^ year^−1^) in the potential total woody stocks (mean and standard deviation) are reported in the bar plots.

**Table 2.** NPP and pCWS computed as average over the simulation period 2006-2099, across all stands and ESMs climate forcing but grouped across RCPs. Mean differences (in percentage) are reported in parenthesis for NPP and pCWS between the alternative management scenarios and the *Business-As-Usual* (BAU) practices used here as the benchmark scenario.

| MANAGEMENT<br>type | RCP | NPP<br>$\text{gC m}^{-2} \text{y}^{-1}$ | pCWS<br>$\text{tC ha}^{-1}$ |
| --- | --- | --- | --- |
| BAU | RCP 2.6 | 501.7 | 192.9 |
| BAU | RCP 4.5 | 522.1 | 195.6 |
| BAU | RCP 6.0 | 530.5 | 196.0 |
| BAU | RCP 8.5 | 542.0 | 198.5 |
| AM+ | RCP 2.6 | 350.0 (−30.2%) | 184.3 (−4.4%) |
| AM+ | RCP 4.5 | 366.8 (−29.7%) | 186.5 (−4.6%) |
| AM+ | RCP 6.0 | 372.0 (−29.8%) | 186.4 (−4.8%) |
| AM+ | RCP 8.5 | 388.1 (−28.4%) | 189.7 (−4.4%) |
| AM− | RCP 2.6 | 495.4 (−1.2%) | 179.8 (−6.6%) |
| AM− | RCP 4.5 | 510.6 (−2.1%) | 181.6 (−7.1%) |
| AM− | RCP 6.0 | 519.9 (−1.9%) | 182.2 (−6.9%) |
| AM− | RCP 8.5 | 524.7 (−3.1%) | 183.8 (−7.3%) |
| NO-MAN | RCP 2.6 | 429.1 (−14.4%) | 136.2 (−29.3%) |
| NO-MAN | RCP 4.5 | 436.4 (−16.4%) | 136.6 (−30.1%) |
| NO-MAN | RCP 6.0 | 444.8 (−16.1%) | 137.1 (−30.0%) |
| NO-MAN | RCP 8.5 | 436.5 (−19.4%) | 136.8 (−31.0%) |

On average, pCWS increased with less intensive management only in the case of a prolonged rotation period under RCP 8.5, while NPP declined (∼6%) in response to an extended rotation and reduced thinning intensity under RCP 8.5. Conversely, pCWS shows the greatest reduction with –13.9% when both thinning intensity and regime only are set to simulate decreased management intensity under RCP 8.5 (Table S5). In summary, AM– simulations generally reduced pCWS with little effect on NPP. For NPP some intra-site differences exist, with the representative forest of Hyytiälä showing higher NPP values when applying a less intensive management than the reference BAU. Differences between the emissions scenarios were small.

### 3.3 More intensive management vs. BAU

Across all stands and ESMs the AM+ simulations reduced NPP about 30% relative to the BAU scenario (Figure 2 and Figure S4, Table 2). Conversely, AM+ reduced pCWS only 4- 5% relative to BAU across the climate change scenarios (Figure 3 and Figure S4, Table 2). (Results for each of the RCP scenarios and under all the alternative management options combined are reported separately across the sites in the Supplementary Material (Table S4 and S5).) The decreases in NPP with AM+ were larger than the decreases with AM- across RCPs, with even a slight increase in NPP in two of the seven AM- scenarios under RCP 2.6 (Table S5). Conversely, pCWS shows a net gain with +4.1%, compared to the BAU, which applies under an increased thinning intensity under RCP 8.5, while the greatest reduction in pCWS with –13.9% is modelled when both thinning intensity and frequency are increased and a shortened rotation turn are simulated under RCP 8.5 (Table S5). Generally, both NPP and pCWS results from the AM+ schemes are, on average, close to or lower than those from the reference BAU and this holds across the three representative forests (Figures S10-12). When considered during the early portion of the study period, the differences in NPP and pCWS between AM+ management scenarios and BAU are narrower. In particular, an average gain of 5% in pCWS could be achieved over a wide range of AM+ schemes, but at the expenses of an average NPP change of –30% (Figure S6).

For both NPP and pCWS, the differences across RCP scenarios are much smaller than across the different management scenarios indicating that management may be more important than near-term climate change in controlling in the C cycle of European forests.

### 3.4 No management vs. BAU

The NPP values in the NO-MAN scenario are, on average, lower than the reference BAU scenario varying from –14.5% (RCP 2.6) to –19.5% (RCP 8.5) (Figure 2, Table 2). However, a site-specific variability in the NPP response to the management scenarios applied exists, with differences between NO-MAN and BAU options ranging from 9.0% to –41.7%, for Hyytiälä and Bílý Křìž respectively, both under the warmest emission scenario (Table S4). Differences between NO-MAN and BAU become more evident during the simulation period and across RCPs scenarios, with the mean NPP value stabilizing or slightly increasing under the BAU (and AM–) option over time. Conversely, in the NO-MAN scenario the values steadily decreased (Figure 2).

The simulated pCWS, which is represented only by standing biomass (i.e., no harvested wood) in the NO-MAN scenario was, on average, lower than in the BAU scenario, with differences of order –30.0% (from –21.6% to –40% across the different sites). During the simulation period, pCWS values in the NO-MAN option first increase slightly at the beginning of the simulation and then decrease significantly toward the end of the century (Figure 3, Table 2). The NO-MAN case returns the lowest average amount of total woody stocks under every emission scenario (Figure 4 and Table 2).

**Figure 4.**
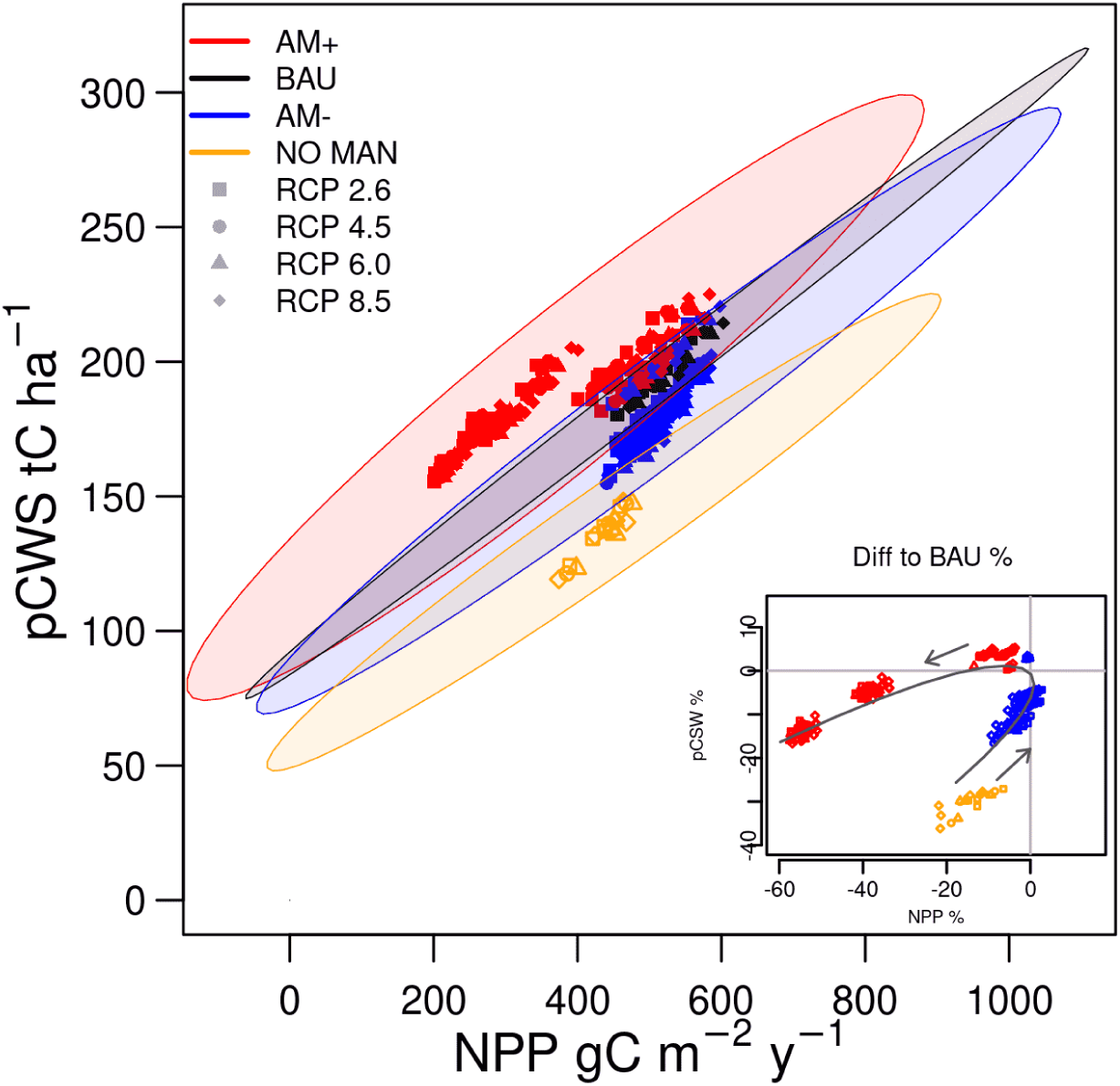
Average NPP (net primary productivity, g C m^−2^ year^−1^) vs. pCWS (the sum of standing and potential harvested woody products; t C ha^−1^) over the period 2006-2099, for the three management scenarios: AM+, AM–, BAU; and the NO-MAN for the 4 RCPs. Reported values refer to data averaged across real and virtual stands and across species. Data ellipses are also reported in shaded colors and refer to all data. NOTE: each single scenario according to Table S1 is reported here (16 in total excluding the mixed ones). In the subplot the differences are expressed as % and are reported along a parametric curve (third order polynomial) with the point (0, 0) representing the reference BAU. Arrows indicate the increasing intensity of management intervention. No significant differences across RCPs were detected.

### 3.5 Mixed management alternatives and the factorial analysis

A mixed combination of management schemes (namely ‘MIX’) was analyzed for all possible combinations (Figure 5 and Supplementary material Table S1, S4 and S5). There were no ‘MIX’ options that simultaneously increased both in NPP and pCWS compared to the BAU scenarios. Values for NPP range from –1.38%, with a prolonged thinning regime and rotation, and an increase in thinning intensity; to –58.4% under a prolonged thinning intensity and rotation period but with a reduced thinning regime, both under RCP 2.6 (Table S5). Similarly, with the same management schemes, pCWS declined 16.1% under RCP 6.0, when compared to BAU, but increased 5.7% under RCP 8.5 in response to increased rotation length in combination with increased thinning intensity.

**Figure 5.**
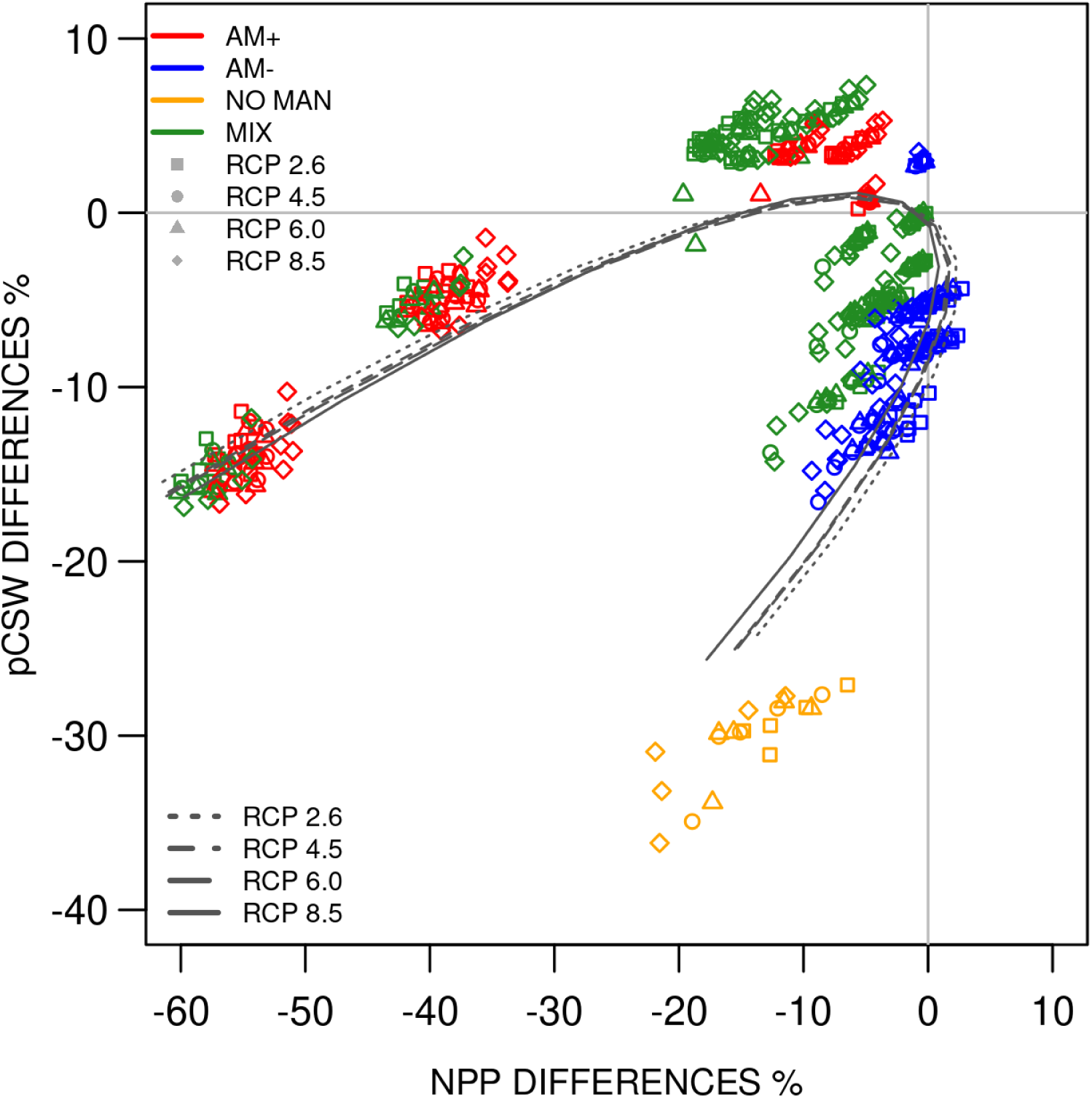
Percentage of changes for Mean NPP (net primary production) and pCWS (potential Carbon Woody Stocks) over the period 2006-2099 between the managed scenarios AM+, AM–, MIX, NO-MAN (AM+, more intensive than BAU; AM–, less intensive than BAU (business as usual); MIX: mixes options according to Table S1, NO-MAN: no management option) and the BAU scenario reported for the four representative concentration pathways (RCPs). Values refer to data averaged across real and virtual stands and across species. Note: each single scenario according to Table S1 is here reported (28 in total). Parametric curve fitting (polynomial of order 3) is also reported for each RCP and it is computed according to the averaged data for each management intervention scenario and RCP combination.

The factorial analysis performed over all the main carbon fluxes and stock variables produced by the model and separated by site, indicates that a significant fraction of the total variability of the key carbon flux variables was driven by the stand factor, i.e. the forest structure as generated by different management schemes, affecting the distribution of age classes, including different above- and below-ground biomass (Figure S6 and Table S6 in Supplementary Material).

## 4. Discussion

### 4.1 Limited leeway to increase carbon uptake and woody stocks with alternative management scenarios

The variables NPP and pCWS represent two sides of the same coin since the former (i.e. NPP) represents the short- to medium-term active carbon sequestration capacity while the latter is the sequestered (and maintained) carbon over the medium- to long-term. The present simulations clearly indicate that even under future climate change scenarios the options are limited for managing forests to help trees maintain their carbon sequestration potential and enhance their ability to cope with increasingly stressful conditions, while in addition increasing their productivity compared to the no-management option. Reducing tree density allows individual trees to benefit from less competition for potentially limiting resources such as light and soil moisture, driving their potential to increase photosynthetic activity and growth rate (Zeide et al., 2001). In addition, stand rejuvenation through harvesting and the replanting of saplings, with less ‘respiring’ (live) biomass per unit of photosynthetic leaf area (because of a shift toward younger stands), and combined with the fertilization effects of increased atmospheric CO_2_ concentration can potentially drives to more productive and efficient forests. This is mirrored in the observed capability of trees to partition more of the photosynthetically assimilated carbon into new woody biomass rather than into nonstructural carbon pools (Campioli et al., 2015; Pappas et al., 2020; Collalti et al., 2020a; Huang et al., 2021). This aspect is reflected in the increasing Harvested Wood Products (HWP; Figure S7) over time with more frequent thinning, reduced tree density, replacement and presence of younger forest stands that potentially can remain in the system as standing biomass over the long-term (Figure S8). Overall, the model projects on average the highest pCWS under the BAU management scheme, even in the future.

The potential to extract more wood and more often, i.e. to shorten the harvest interval, and at the same time maintain at least the current forest biomass, depends on NPP under the different scenarios. We found, however, that BAU management remains the most favorable scheme under future environmental conditions and might already be a close-to-optimum management approach for different RCP scenarios (Figure 4) and across the individual sites (Figures S9-S11). This is an endorsement of past research arriving at today’s management practices and a coincidence that today’s most favorable scheme might also be most favorable in a future altered climate. With more frequent harvesting and replanting and increasing intensity of intervention compared to the benchmark BAU, the NPP is not shown to increase any further under any RCP scenario, in spite of an average younger and, in theory, more productive forest stand. The net growth rate does not compensate for the increased fellings (tree harvesting), while in parallel there is a limited yield in terms of increased woody carbon stocks, as reflected in a low standing biomass that is likely a sign of a critically low tree density. Albeit the BAU reference benchmark is already an intensive management approach with tree fellings as a percentage of net annual increment of 84%, 77% and 101% for Czech Republic, Finland and Denmark, respectively, as reported for 2005 (State of Europe’s forests 2020), the first year of our RCP-based climate change response simulations. Similarly, Pussinen et al. (2009) showed that increasing the total harvested products led to a decrease in both NPP and forest standing biomass in some European areas. The difficulties associated with simultaneously increasing both forest standing biomass and harvested wood products were shown in the seminal modeling study of Thornley and Cannell (2000).

An important factor contributing to the apparent lack of significant differences in forest responses across RCPs scenarios, compared to the differences across different management schemes, might come from the combination of counteracting key drivers of plant physiology (e.g. lengthening of the growing season by warming and, in parallel, an increased maintenance respiration rate from that same warming) which are considered in the model despite temperature acclimation. Although experimental evidence for the CO_2_ fertilization effect on plant carbon accumulation is strong and is typically predicted by vegetation models albeit with different degrees of certainty, the probability for its persistence into the longer-term future is a hotly debated issue (Nabuurs et al., 2013; Habau et al., 2020; Wang et al., 2020; Gatti et al., 2021; Walker et al., 2021). The biochemical model of photosynthesis used here (Farquhar et al., 1980) includes a nonlinear saturating response to CO_2_, yet, other environmental drivers, such as temperature, vapor pressure deficit (which scales exponentially with warming) and water availability were shown to interact to down-regulate the positive CO_2_ effect on GPP (Grossiord et al., 2020). Data at the biome scale (see Luyssaert et al., 2007) indicated a potentially high sensitivity of plant respiration to warming that may stabilize NPP over a temperature threshold with no further gains. Warming in low-temperature-limited forest biomes would be expected instead to have a positive effect on annual GPP and NPP (Henttonen et al., 2017; Sedmáková et al., 2019). However a warming-induced increased respiration cost might curb these trends and even offset a positive GPP and/or NPP response to increasing atmospheric CO_2_ concentration, as indicated by some other modeling and experimental studies (Way et al., 2008; Gustavson et al., 2017; Collalti et al., 2018; but see Reich et al., 2016). For example, Mathias and Trugman (2021) showed a potential future unsustainable growth for boreal and temperate broadleaved forests, with the net overall effect of decreased NPP. Other studies already indicated that combined impacts of warming and increasing atmospheric CO_2_ concentration might cause forests to grow faster and mature earlier but also to die younger (Kirschbaum, 2005; Collalti et al., 2018, 2019). With the increasing standing biomass and accumulation of more respiring tissue in older trees, plant respiration might increase more quickly than GPP, as the canopy closure would be reached earlier, capping GPP when all available solar radiation is intercepted, but with sustained respiratory needs. The use here of many virtual stands of different ages in our simulations might have (realistically) compensated for local stand-age effects on biomass and stand structural responses to climate change that would be counteracting across a landscape. To the extent that this is true, the simulated patterns obtained would be related to effects of climate, forest management, and their multiple combinations only.

Ultimately these simulations indicate that increasing the harvest rate and at the same time keeping the fellings below the growth rate will be challenging. As such, the possibility of simultaneously increasing both carbon sequestration rate and tree carbon (standing biomass) storage capacity while managing forests in a sustainable way may be very limited. A steady intensification or intervention frequency – alone or in combination – compared to the business-as-usual scheme might come at the price of a substantial loss of primary productivity. While the amount of potential harvested woody products still would be significant, we would *de facto* end up reducing the active forest carbon sink and thus the forest’s potential to assimilate and sequester CO_2_ from the atmosphere.

### 4.2 Role of forest management in the context of climate change

In the context of climate uncertainty and because of policy intentions, management practices may no longer prioritize productivity only – which traditionally includes rotation times being adjusted to maximize value of timber – without preserving the forest carbon sink and ensuring the long-term functioning of forests and the continued provision of their many ecosystem services (Krofcheck et al., 2019). The selection of adaptive management practices has been suggested as a mechanism to potentially enhance the climate change mitigation potential of forest ecosystems (Tahvonen, 2016; Yousefpour et al., 2017). Our model results highlight, for Central and Northern temperate and boreal European forests, the importance of forest structure to sustain or even enhance productivity and carbon storage. This in turn indicates that management practices may be quantitatively as important as future climate and atmospheric CO_2_ concentration in regulating the carbon sink strength of forests, which is in line with some previous modeling studies (Garcia-Gonzalo et al., 2007; Akujärvi et al., 2019; de Wergifosse et al., 2022). These simulations indicate that silvicultural practices included in the model would persist as key factors in the regulation of carbon sequestration through the end of this century under any of the CMIP5 RCP scenarios. In accordance with the modeling study of Kindermann et al. (2013), our results indicate the need to sustain NPP rather than maximizing forest carbon stocks. Our results also point out, however, a narrow operational space surrounding the BAU scheme which can be designated as near-optimal over a wide and diversified portfolio of alternative management schemes across the broad range of RCP/ESM-based climate change scenarios. Conversely, other studies (Garcia-Gonzalo et al., 2007; Luyssaert et al., 2018) showed that harvest intensity should be loosened in order to maximize the carbon sink which would lead to reductions in the wood harvesting rates. However, under adverse climate change effects, reductions in wood harvesting rates may correspond with declining carbon sequestration.

Even considering changes in species composition by replanting better-suited species under a climate change adaptive framework, Schelhaas et al. (2015) found a reduction of the net increments, although these did not affect wood product amounts. On the other hand, Pussinen et al. (2009), found that under an increased felling rates regime (i.e. high harvested products amount) maintaining the current forest standing biomass was possible even under future climate scenarios. However, in their study NPP was simply assumed to increase proportionally to the temperature, driving increased growth rates at boreal and temperate forest sites. The present simulation study used a more realistic process-based representation of climate change impact on forest growth. This reveals instead a more modest, or even unfavorable, effect of climate change despite a CO_2_ fertilization effect on NPP. Others have suggested that past and/or future climate change did, or could, negatively affect NPP (Reich and Oleksyn, 2008; Bastos et al., 2020) in a range of forested and non-forested ecosystems through increased frequency and/or magnitude of large-scale disturbances (e.g. heat waves, windstorms, weather-based pest outbreaks), with significant variation in effects in different ecosystems or forest types and locations (e.g. Thom et al., 2017; Nabuurs et al., 2019; McDowell et al., 2020; Senf and Seidl, 2021b; Gampe et al., 2021). Should such increases in climatic extremes (e.g. drought and heatwaves) and disturbances (pests outbreaks, wildfires, storms) occur -- and negatively affect the carbon sink capacity of a significant fraction of European forests -- the robustness of the BAU management scheme specified in our simulations might be questioned with an adaptive set of management options designed, tested and ultimately put into place (Yousefpour et al., 2017). However, under the unmanaged- forest scenario, our simulations resulted in significant and steady decline in NPP through the year 2099 when averaged over all the climate change scenarios considered.

### 4.3 Further consideration and outlook

In this study we found that intensification of forest management practices might lead to declines in both CO_2_ assimilation and the total woody stocks.

The focus was on a subset of forest types that currently play a significant role in the European carbon balance and wood product markets by representing more than 50% of the EU standing biomass (Avitabile et al., 2020).

In the current modelling framework, our results refer to potential values of carbon woody biomass, with no estimated decay for harvested wood products.

In this context, our aim was primarily to quantify the efficiency of forest trees to actively sequester and store carbon from the atmosphere and, thus, defining their potential contribution to mitigation policy requirements (see e.g. Valade et al., 2017).

Pure stands were selected for this study as they represent most of the managed stands in Europe, and no species migration under climate change was included, even in the no- management scenario. However, we highlight that the overall climate change mitigation of the forestry sector is further influenced by carbon sequestration in harvested wood products (Brunet-Navarro et al., 2016) as well as energy and material substitution effects of wood products, which is depending on dynamic changes in energy portfolios and greenhouse gas emissions in other sectors and thus associated with substantial and additional uncertainty (e.g. Leskinen et al., 2018; Howard et al., 2021; Soimakallio et al., 2021).

Since the rates of potential species migration and/or replacement may be incompatible with the expected rates of climate change, at least for the high RCP scenarios (Settele et al., 2014), the effectiveness of tree species change as a mitigation factor in unmanaged forests could be limited. Future research with validated dynamic vegetation models is paramount for it might provide needed insights into European forest carbon sequestration potential across a broad geographic extent through mechanisms of seed dispersal, species migration and up-scaling techniques (Fritsch et al., 2020).

In addition, despite simulating a broad and diversified range of different silvicultural interventions aimed at testing the possibility to increase NPP and carbon stocks, additional studies will be crucial to address changes in species selection (i.e. genetic selection) and assisted species migration in adaptive forest management.

Finally, the focus of the present work was to quantify the capacity of trees to sequester carbon in woody biomass and in turn to provide wood products in the face of expected climate changes. A life-cycle analysis of the harvested wood products was not included, and in essence implicitly assumed to be unaffected by climate change.

## 5. Conclusions

To our knowledge this is one of the first modeling studies that systematically analyses, over a wide range of scenarios, the possibilities and limitations of altering forest management practices to achieve the twofold objective of maximizing forest C-sequestration capacity while concomitantly maintaining and/or increasing pCWS in the face of future climate change.

Even though the analysis was confined to three sites, the representativeness of those sites to other European forests and the general consistency of the results indicates a broader significance and applicability. In particular, the differences between AM+ and AM–, each relative to BAU, indicates a relatively consistent response across species within different climate change scenarios and management options.

Our results indicate that the scope to meet the above twofold objective may be limited, because business-as-usual management practices may already be nearly optimal in terms of carbon uptake, sequestration and storage, as testament to positive outcomes of previous silvicultural research. A general conclusion of the modelled results is that NPP and/or pCWS are likely to decline, relative to BAU, with any significant change in forest management practices.

Beside the economic value of the extractable wood and the potential for energy and material substitution, it is today crucial for EU countries to preserve forest’s functionalities under the pressure of the rapidly changing climate conditions, in order to maintain the climate mitigation potential and the supply of wood products and many ecological goods and services. Forest management based on scientific principles remains a valuable tool for local, regional and global strategies to optimize the forest carbon sinks and provide desired products under a varying climate. However, the extensive modeling in this study and related research does not support an optimistic view that changes to management practices in European forests can be relied on to significantly increase carbon uptake and storage while increasing wood production above current rates.

## Acknowledgements

We are thankful to A. Ibrom, P. Kolari and J. Krejza for providing us data for Sorø, Hyytiälä and Bílý Kříž sites. We thank J. Holder, B. Waring, H. Bugmann, L. Perugini, S. Luyssaert, M. Michetti, and A. Mäkelä for comments on a previous version of the manuscript and/or fruitful discussions. We also thank the ISI-MIP project (https://www.isimip.org/) and the COST Action FP1304 PROFOUND (Towards Robust Projections of European Forests under Climate Change), supported by COST (European Cooperation in Science and Technology) for providing us the climate historical scenarios and site data used in this work. This work used eddy covariance data acquired and shared by the FLUXNET community, including these networks: AmeriFlux, AfriFlux, AsiaFlux, CarboAfrica, CarboEurope-IP, CarboItaly, CarboMont, ChinaFlux, Fluxnet-Canada, GreenGrass, ICOS, KoFlux, LBA, NECC, OzFlux-TERN, TCOS-Siberia, and USCCC. The ERA-Interim reanalysis data are provided by ECMWF and processed by LSCE. The FLUXNET eddy covariance data processing and harmonization was carried out by the European Fluxes Database Cluster, AmeriFlux Management Project, and Fluxdata Project of FLUXNET, with the support of CDIAC and ICOS Ecosystem Thematic Center, and the OzFlux, ChinaFlux, and AsiaFlux offices. We acknowledge the World Climate Research Programme’s Working Group on Coupled Modelling, which is responsible for CMIP, and we thank the respective climate modeling groups for producing and making available their model output. The U.S. Department of Energy’s Program for Climate Model Diagnosis and Intercomparison at Lawrence Livermore National Laboratory provides coordinating support for CMIP and led development of software infrastructure in partnership with the Global Organization for Earth System Science Portals. A.C. was supported by the EFI-Funded FORMASAM (FORest MAnagement Scenarios for Adaptation and Mitigation) project; D.D. and A.C. were also supported by resources available from the Italian Ministry of University and Research (FOE-2019), under projects ‘Climate Changes’ (CNR DTA. AD003.474.029); A.C. and G.M. were supported by LANDSUPPORT Horizon 2020 research and innovation programme under grant agreement No. 774234. The 3D-CMCC-FEM model code is publicly available and can be found on the GitHub platform at: https://github.com/Forest-Modelling-Lab/3D-CMCC-FEM). All data supporting this study are publicly available at the Zenodo repository (https://zenodo.org/record/6026117#.YgTfXnXMJhE). Correspondence and requests for additional materials should be addressed to corresponding authors.

## Credit authorship contribution statement

D. Dalmonech, G. Marano and A. Collalti performed conceptualization, data curation, formal analysis, investigation, writing the original draft, and editing; C. Trotta ran the model code; J. Amthor, M. Lindner and A. Cescatti, supported for writing, reviewing and editing.

## Supplementary Material

**Figure S1.**
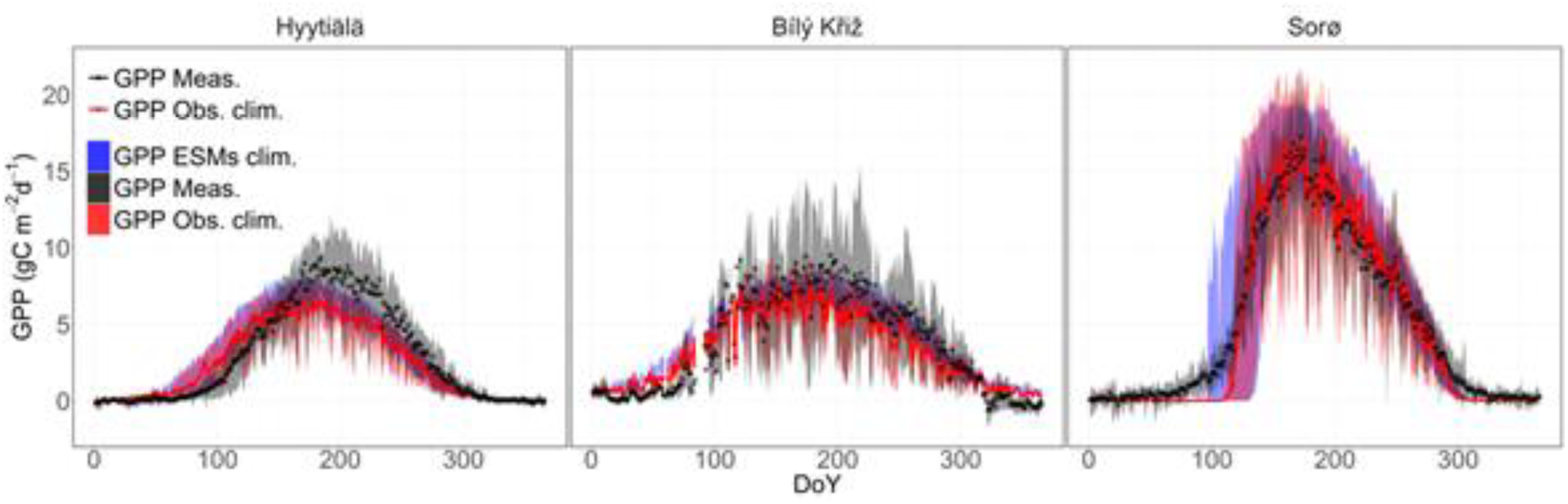
Evaluation of seasonal GPP (gross primary productivity) fluxes. Top row shows the seasonal course of average daily GPP (g C m^−2^ day^−1^) over the years (DoY = Day of the Year). Blue shaded area represents the maximum and minimum bounds of GPP values among the five ESMs used to force the model (GPP ESM clim.), red line and the red shaded area represent the average and the maximum and minimum GPP values when the model is forced by observed climate (GPP Obs. clim.), and black dots represent the quality-checked and - filtered GPP values evaluated at the sites by the eddy covariance technique (GPP Meas.) and the shaded area of the inter-annual variability. ESMs = model forced with Earth System Models climate. Performance statistics are reported in Table S2.

**Figure S2.**
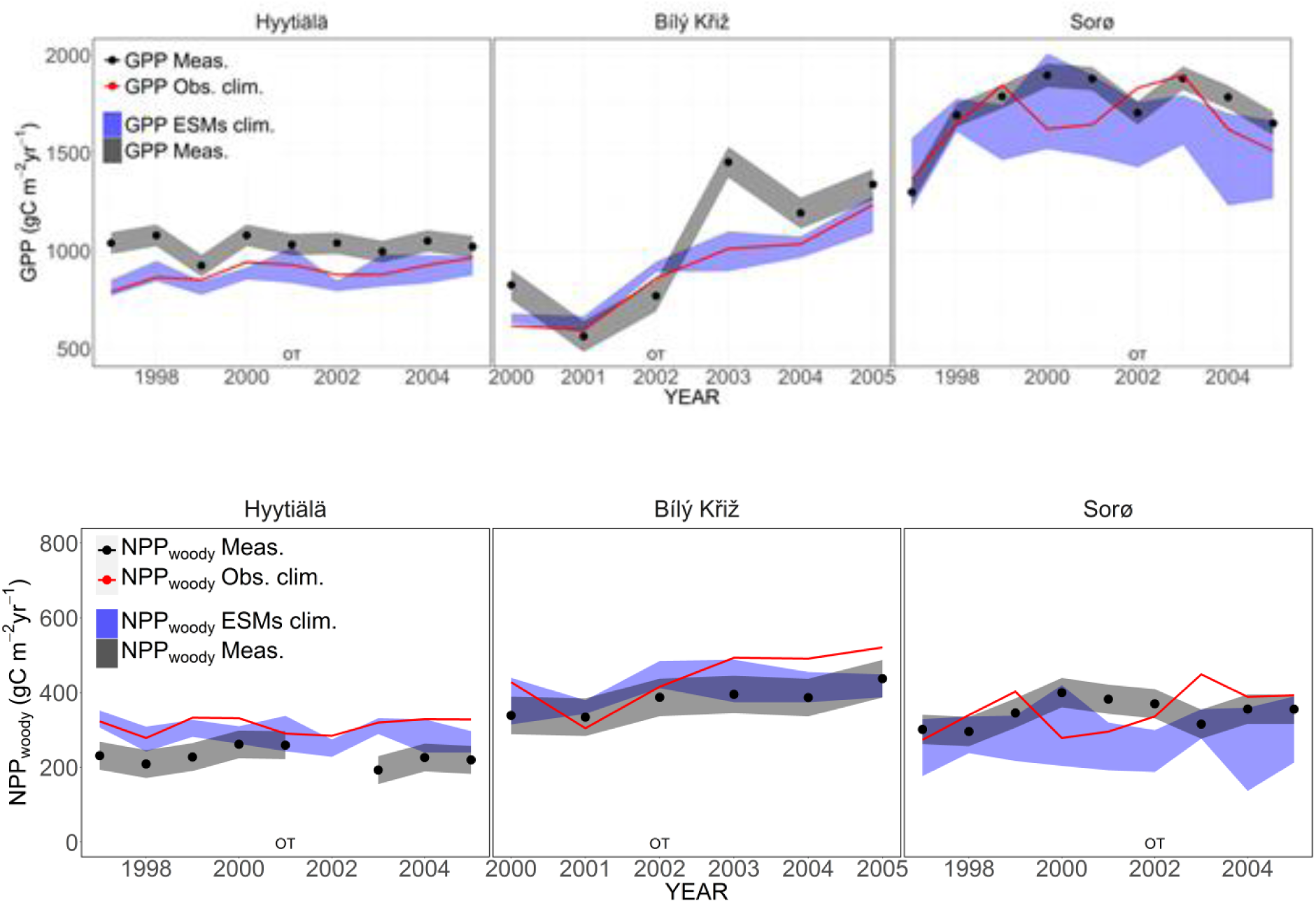
Comparison of the predicted annual GPP (gross primary productivity, gC m^−2^ day^−1^, above panel) and NPP_woody_ (the annual NPP allocated to woody pools, gC m^−2^ day^−^1, below panel) with site observations at the three sites modelling real stands. The blue shaded area represents the maximum and minimum bounds of GPP and NPP_woody_ values among the five ESMs used to force the model; the red line represents the average NPP_woody_ values when the model is forced by observed climate. Black dots represent the measured GPP and NPP_woody_ values (i.e. GPP Meas. and NPP_woody_ Meas.), and the gray shaded area represents the relative uncertainty bounds. At the Hyytiälä site NPP_woody_ observed data for the year 2002 was missing. OT = observed thinning at the site; ESM = model forced with Earth System Models climate.

**Figure S3.**
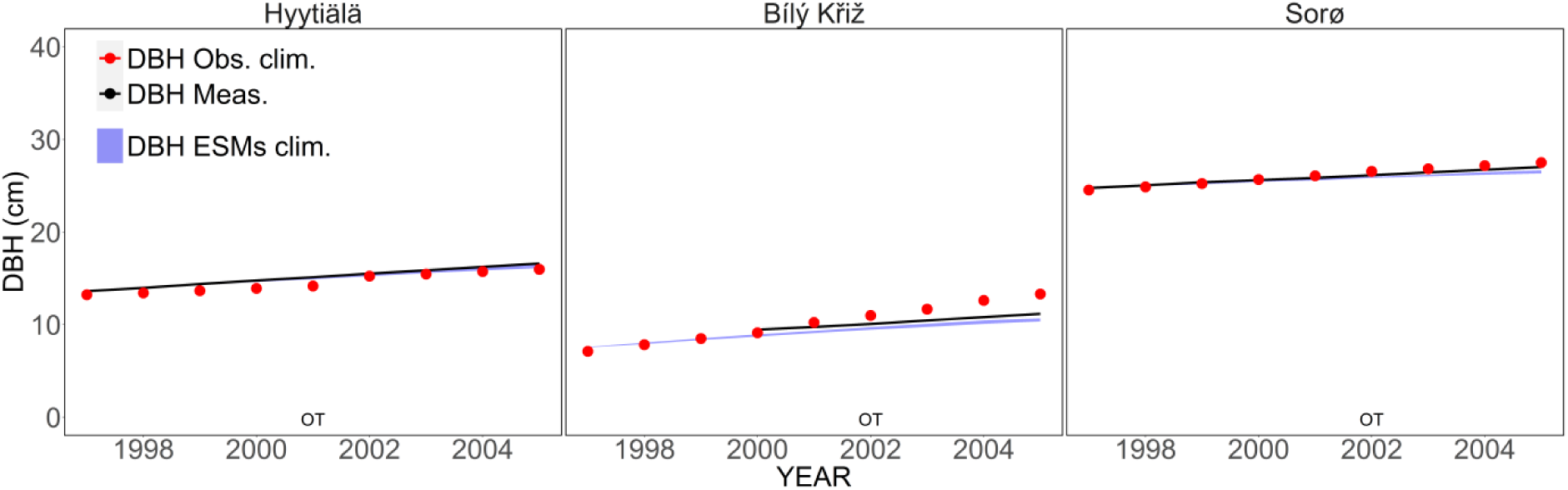
Comparison of the predicted annual DBH (stem diameter at breast height in cm) with site observations at the three sites. The shaded area represents the maximum and minimum bounds of DBH values among the five ESMs used to force the model; the red line represents the average DBH values when the model is forced by observed climate (DBH Obs. clim.). Black dots represent the measured DBH values. OT = observed thinning at the sites; ESMs = model forced with Earth System Models climate.

**Figure S4.**
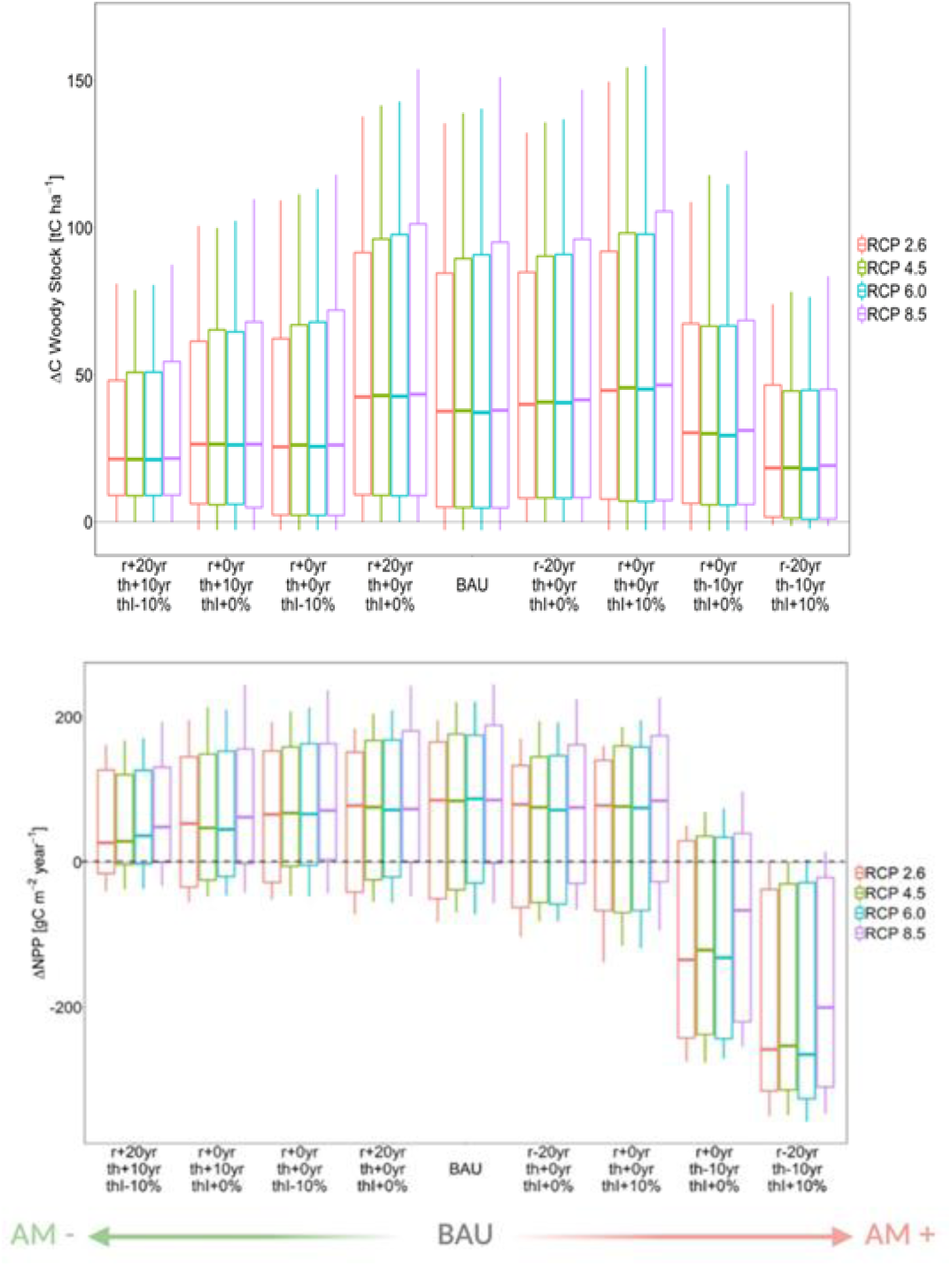
Box plots of the differences for pCWS (potential C Woody Stocks), (a) computed with respect to the unmanaged scenario NO-MAN (dashed line) under different RCPs and NPP (net primary production) (b). Management scenarios are reported according to changes in rotation turn (*r*), thinning frequency (*th*) and thinning intensity (*thi*) and according to the AM+ scenarios (more intensive than BAU) and AM– (less intensive than BAU) reported in Table S1. The management key variables are changed with respect to the BAU scenario (r+20 th+10 thi –10, r+00 th+10 thi 00, r+00 th+00 thi –10, r+20 th+00 thi+00 belongs to the less intensive scenario AM–; r–20 th+00 thi+00, r+00 th+00 thi+10, r+00 th–10 thi+00, r–20 th–10 thi+10 belongs to the more intensive scenario AM+). Scenarios from the BAU are ordered following the most important management variable driving the fluxes and stocks variability, i.e. thinning frequency and rotation (data not shown). For the sake of clarity we reported 8 out of 14 scenarios in the figure. However, the main pattern across intensities does not change.

**Figure S5.**
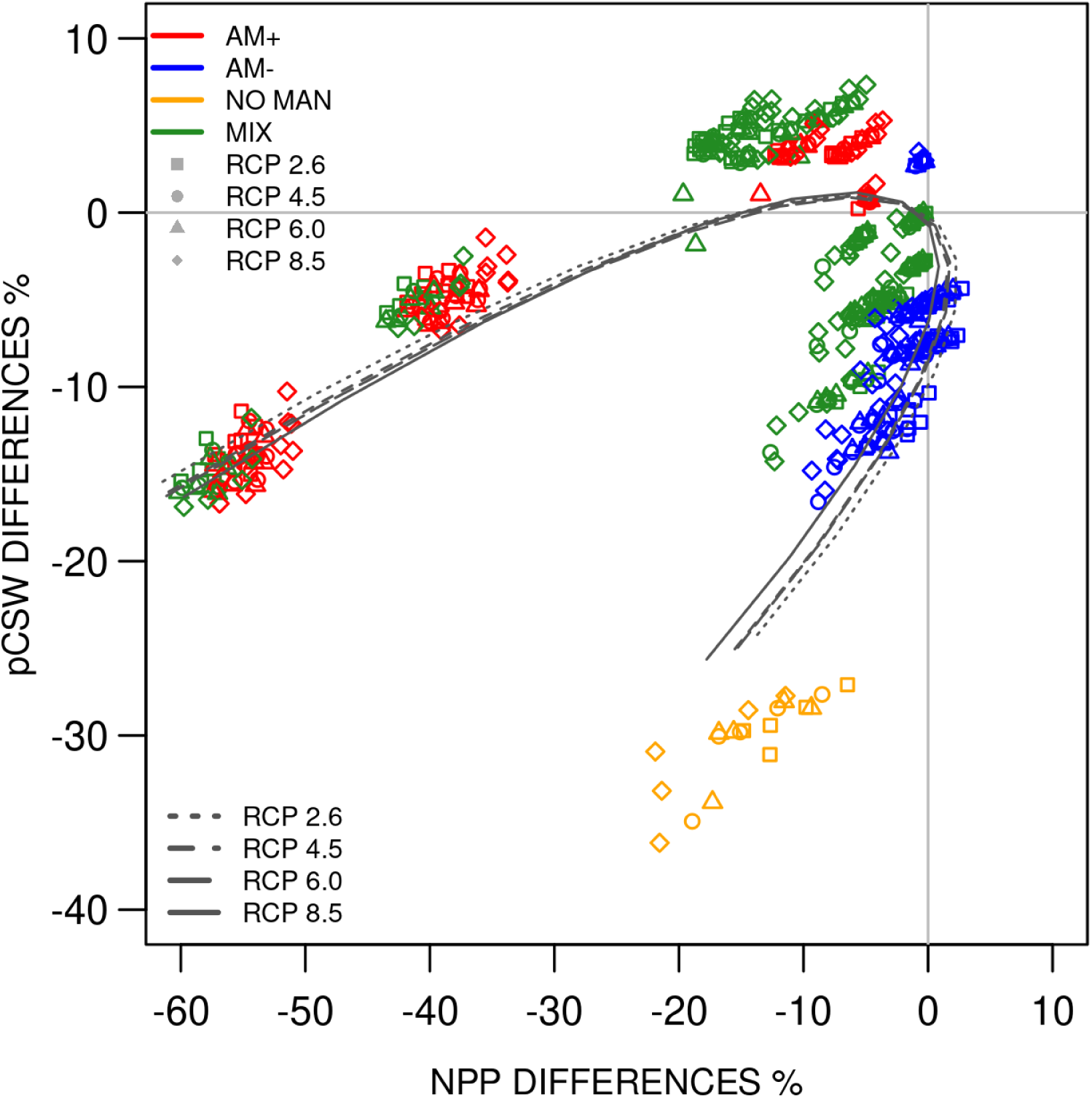
Percentage of changes for Mean NPP (net primary productivity) and pCWS (potential Carbon Woody Stocks) over the period 2006-2099 between the managed scenarios AM+, AM–, MIX, NO-MAN (AM+, more intensive than BAU; AM–, less intensive than BAU; MIX: mixes options according to Table S1, NO-MAN: no management option) and the BAU (business–as–usual) scenario reported for the 4 RCPs. Values refer to data averaged across real and virtual stands and across species. Note: each single scenario according to Table S1 is here reported (28 in total). Parametric curve fitting (Polynomial of order 3) is also reported for each RCP and it is computed according to the averaged data for each management intervention scenario and RCP.

**Figure S6.**
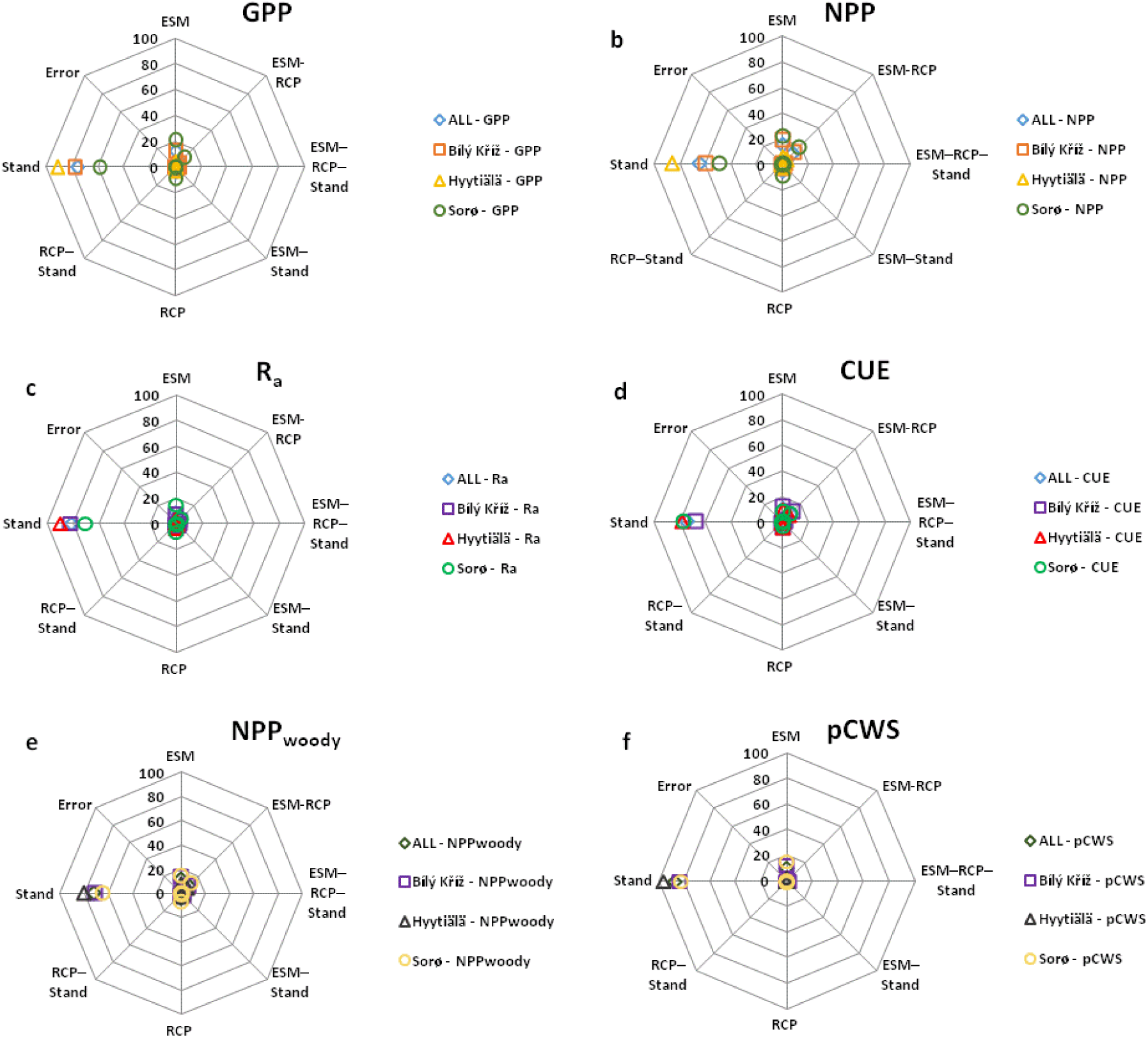
Factorial analysis of the factors driving the variability of GPP, NPP, R_a_, CUE, NPP_woody_ and pCWS, for the three study sites Bílý Křìž (square), Hyytiälä (triangle), Sorø (circle) and an average among sites (rhombus); ESMs = model forced with Earth System Models climate, RCP = emission scenarios, ‘Stand’ = virtual stands of the Composite Forest Matrix, i.e. simulations from each individual virtual stand, characterized by a specific age class and structure, are considered.

**Figure S7.**
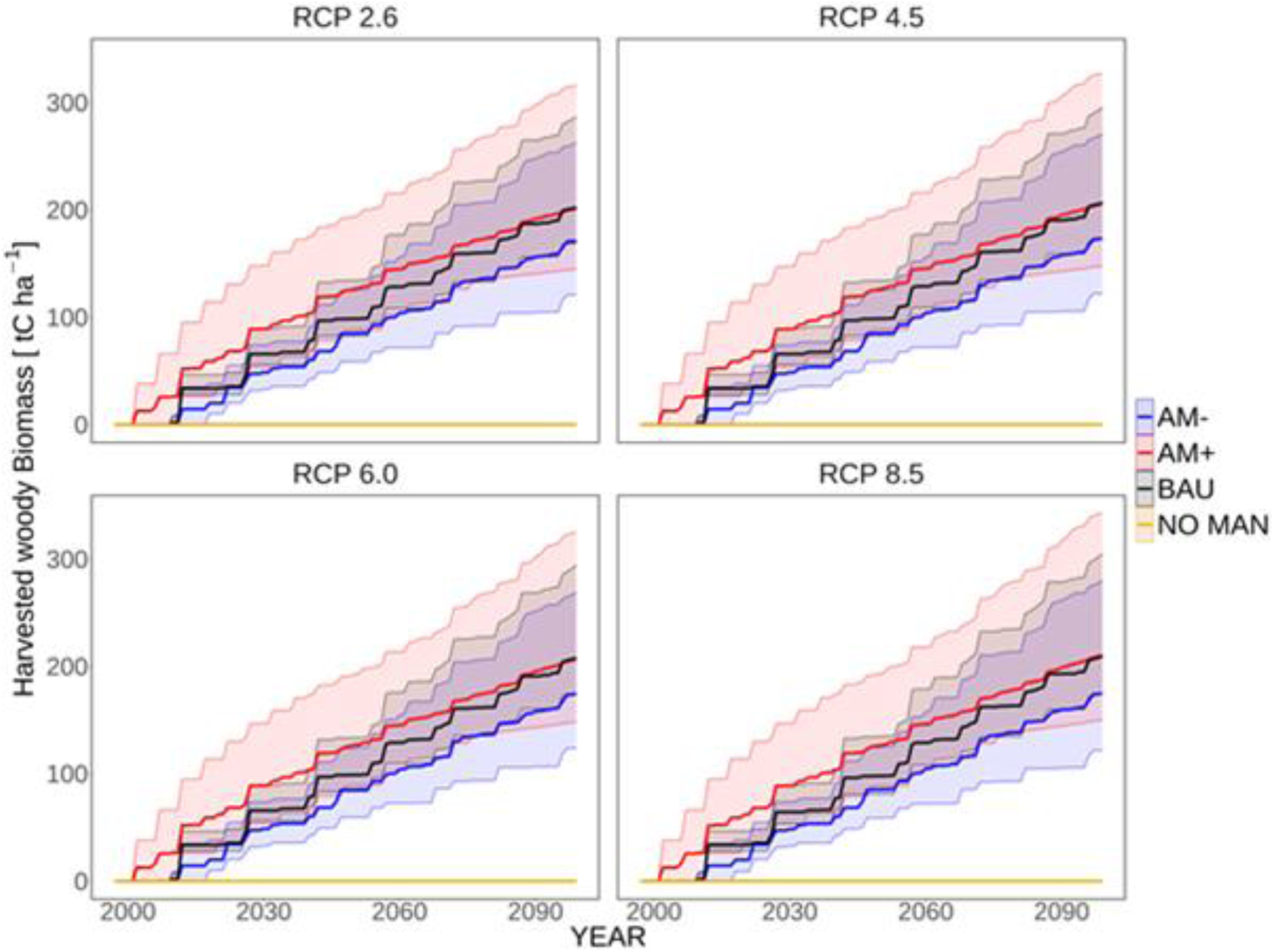
Potential harvested woody Biomass (pHPW) simulations under the management scenarios (AM+, BAU, AM–) and the NO-MAN scenario divided by different emission scenario RCPs. pHWP, solid line, is averaged across the representative forests, different ESMs and aggregated according to the management regime. Yearly mean values and 5^th^ and 95^th^ percentiles are reported.

**Figure S8.**
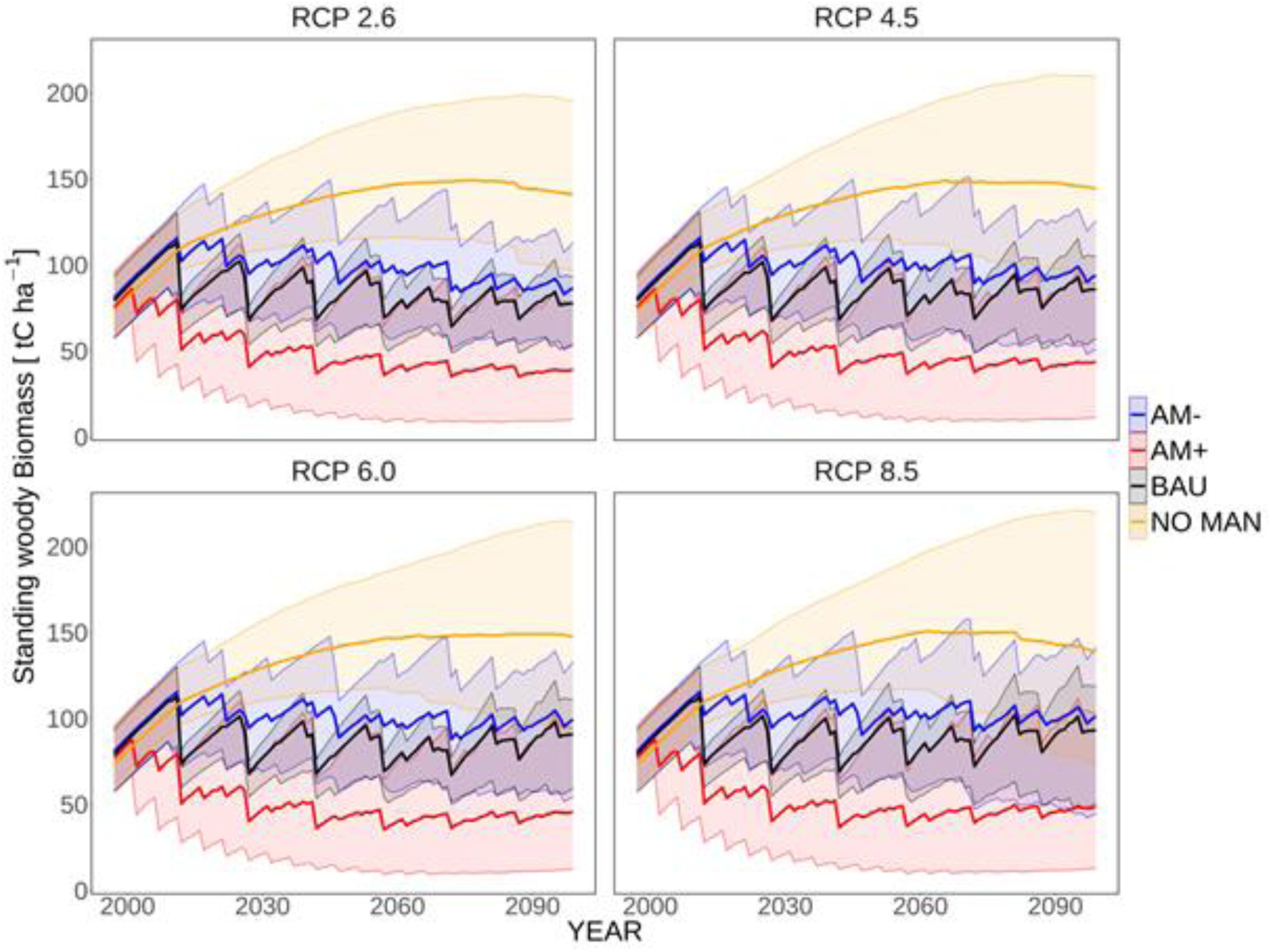
Standing woody biomass simulations under the management scenarios (AM+, BAU, AM–) and the NO-MAN scenario divided by different emission scenario RCPs. Standing woody biomass, solid line, is averaged across the representative forests, different ESMs and aggregated according to the management regime. Yearly mean values and 5^th^ and 95^th^ percentiles are reported.

**Figure S9.**
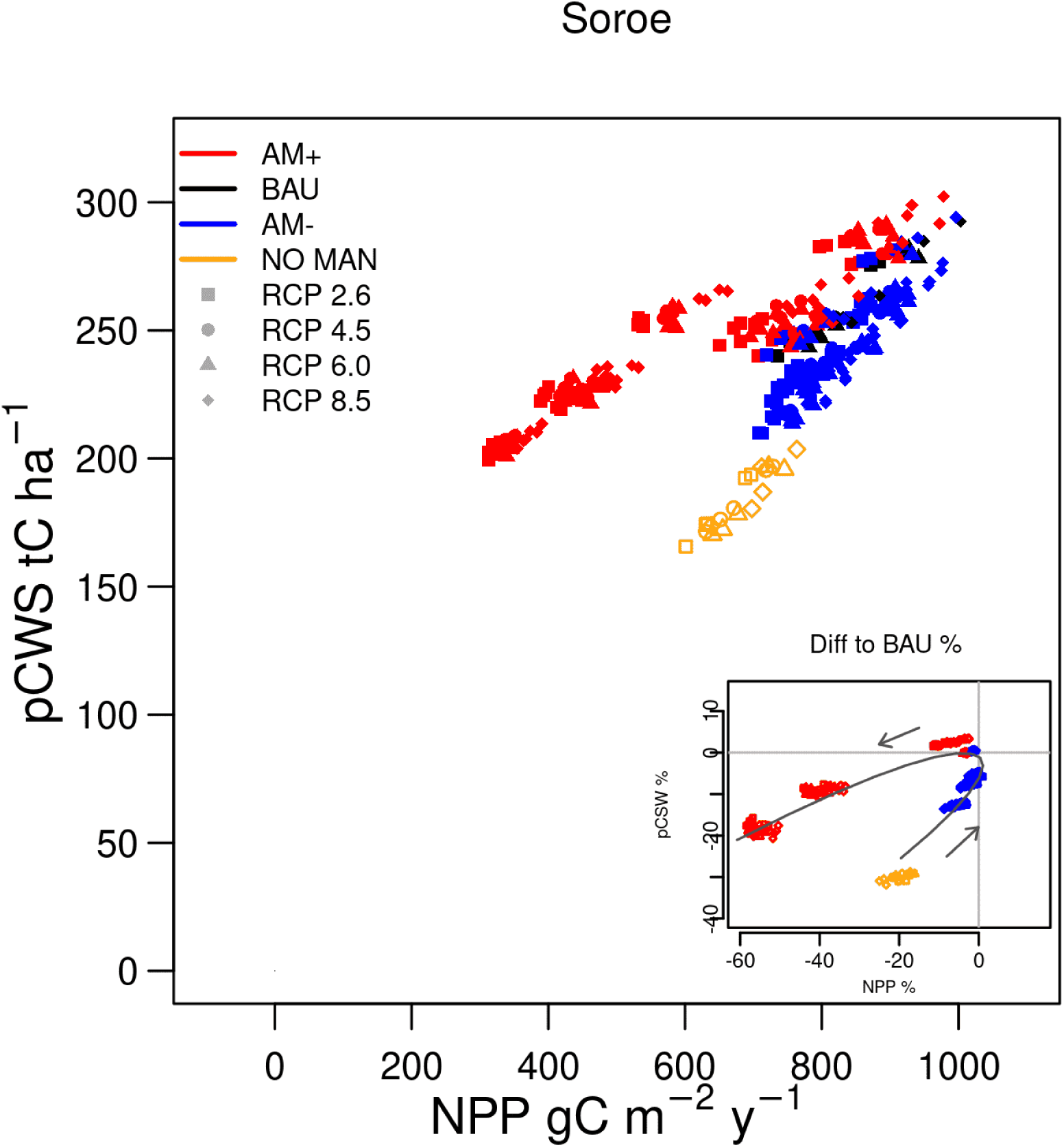
Average NPP (net primary productivity, g C m^−2^ year^−1^) and pCWS (the sum of standing and potential harvested woody products; t C ha^−1^) over the period 2006-2099, for the three groups of management scenarios: AM+, AM–, BAU; and the NO-MAN reported for the 4 RCPs at the site of Sorø. Reported values refer to data averaged across real and virtual stands. Data ellipses are also reported in shaded colors and refer to all data. NOTE: each single scenario according to Table S1 is reported here (16 in total). In the subplot the differences are expressed as % and are reported along a parametric curve (third order polynomial) with the point (0, 0) representing the reference BAU. Arrows indicate the increasing intensity of management intervention.

**Figure S10.**
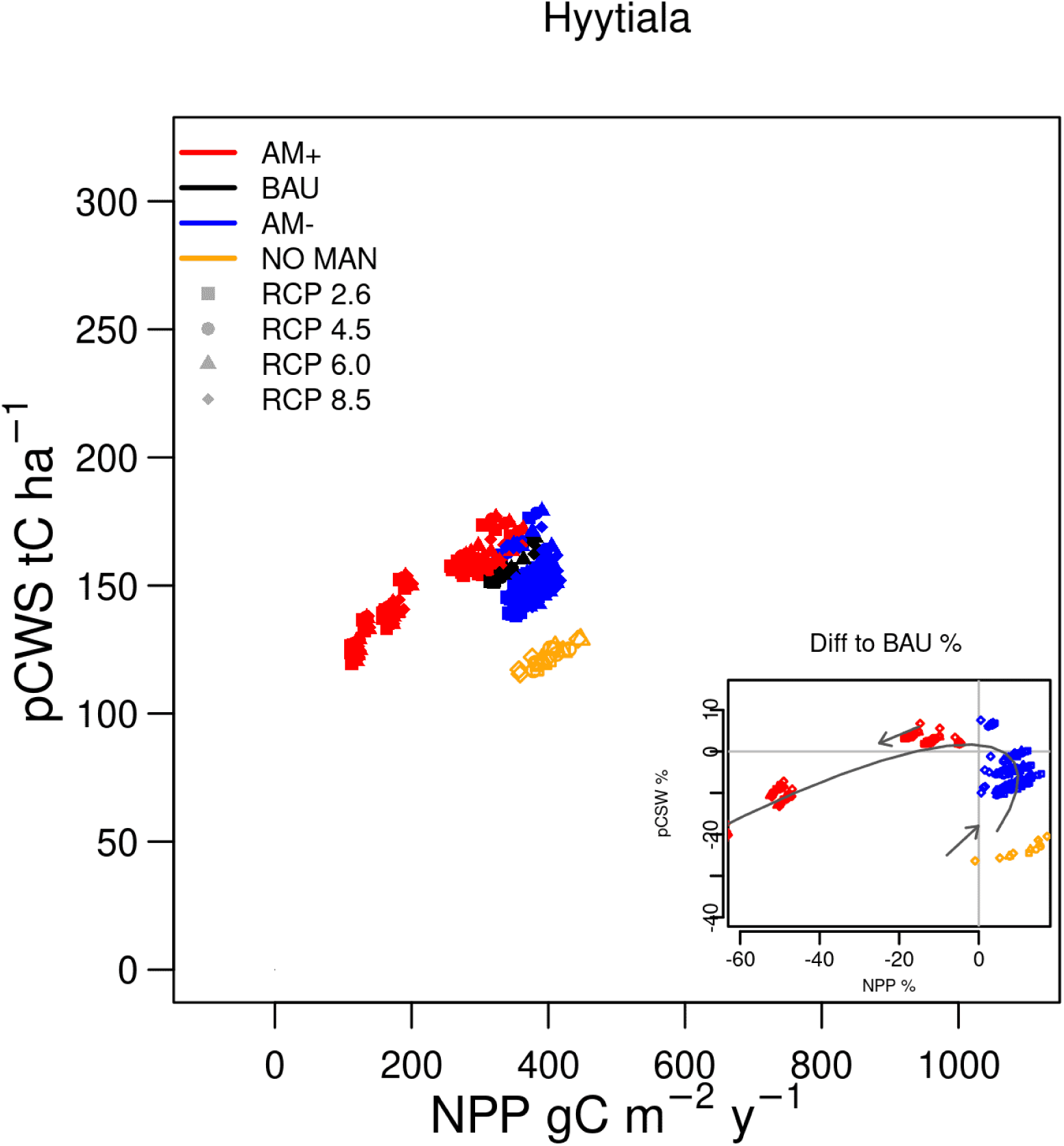
Average NPP (net primary productivity, g C m^−2^ year^−1^) and pCWS (the sum of standing and potential harvested woody products; t C ha^−1^) over the period 2006-2099, for the three groups of management scenarios: AM+, AM–, BAU; and the NO-MAN reported for the 4 RCPs at the site of Hyytiälä. Reported values refer to data averaged across real and virtual stands. Data ellipses are also reported in shaded colors and refer to all data. NOTE: each single scenario according to Table S1 is reported here (16 in total). In the subplot the differences are expressed as % and are reported along a parametric curve (third order polynomial) with the point (0, 0) representing the reference BAU. Arrows indicate the increasing intensity of management intervention.

**Figure S11.**
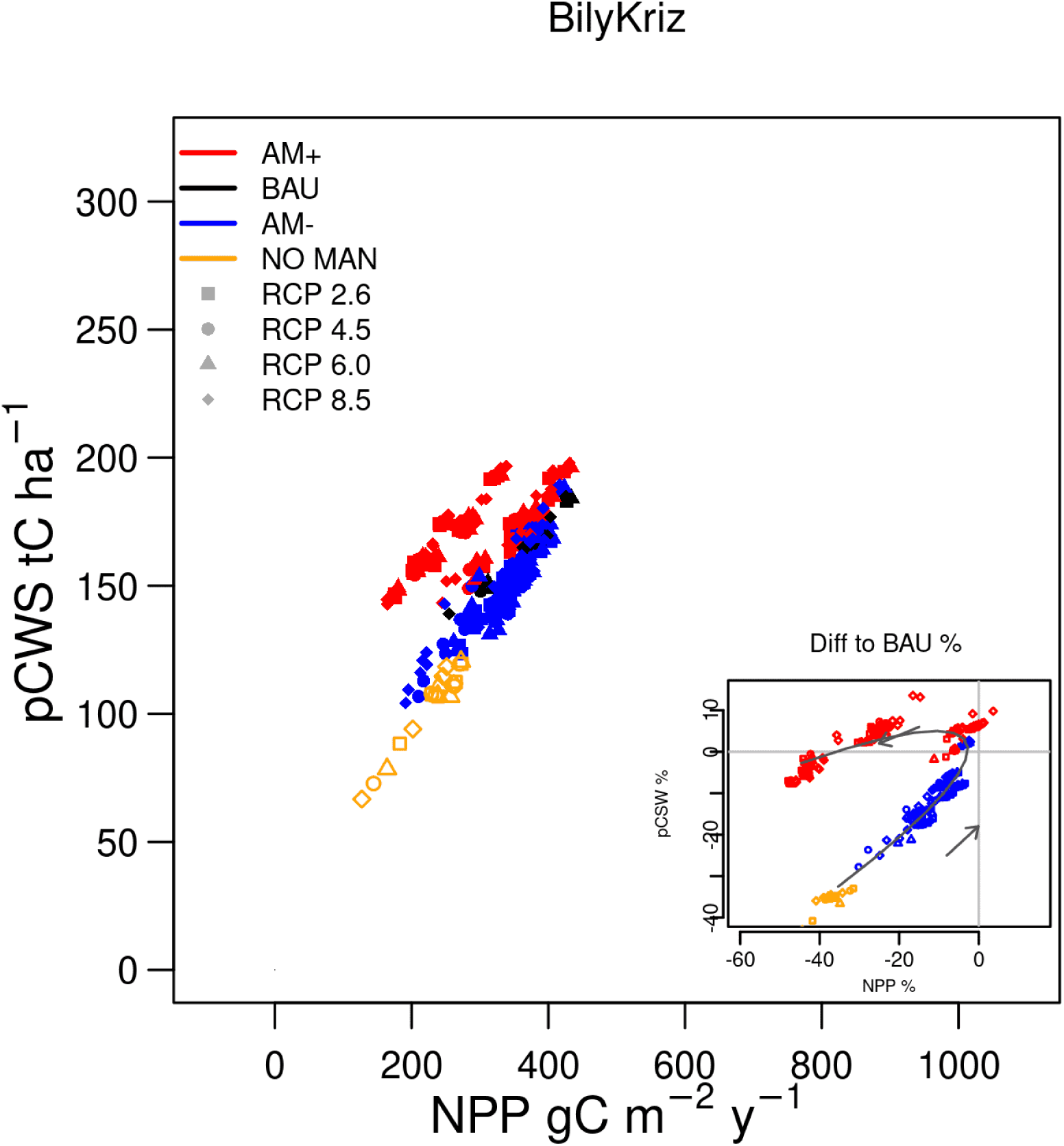
Average NPP (net primary productivity, g C m^−2^ year^−1^) and pCWS (the sum of standing and potential harvested woody products; tC ha^−1^) over the period 2006-2099, for the three groups of management scenarios: AM+, AM–, BAU; and the NO-MAN reported for the 4 RCPs at the site of Bílý Křìž. Reported values refer to data averaged across real and virtual stands. Data ellipses are also reported in shaded colors and refer to all data. NOTE: each single scenario according to Table S1 is reported here (16 in total). In the subplot the differences are expressed as % and are reported along a parametric curve (third order polynomial) with the point (0, 0) representing the reference BAU. Arrows indicate the increasing intensity of management intervention.

### Supp. Tables

**Table S1.** Full experimental design of management scenarios: 26 alternative scenarios, plus an unmanaged scheme (NO-MAN) where trees are removed only through mortality, each relative to the business-as-usual scheme. Scenarios ranged from more limited interventions, grouped under the label AM–, to more intensive silvicultural practice, grouped under the label AM+. The cutting scenarios are a combination of three different thinning intensities at three different thinning intervals and three different rotation ages (e.g. rotation = +20, thinning intensity = 10, thinning regime = –10 indicate increasing the rotation period by 20 years with respect to the BAU setting, increasing the thinning interval by 10 years relative to the BAU setting, and decreasing the intensity of removal by 10% relative to the BAU setting (Table 1). A mixed combination of scenarios, grouped under the label MIX, was also simulated and indicated where the parameter changes follow opposite directions (e.g. increase thinning intensity in combination with prolonged rotation period).

| Management type | rotation years | thinning regime years | thinning intensity % |
| --- | --- | --- | --- |
| AM- | 20 | 10 | -10 |
| AM- | 0 | 10 | -10 |
| AM- | 0 | 10 | 0 |
| AM- | 20 | 10 | 0 |
| AM- | 20 | 0 | -10 |
| AM- | 0 | 0 | -10 |
| AM- | 20 | 0 | 0 |
| BAU | 0 | 0 | 0 |
| AM+ | -20 | 0 | 0 |
| AM+ | 0 | 0 | 10 |
| AM+ | -20 | 0 | 10 |
| AM+ | -20 | -10 | 0 |
| AM+ | 0 | -10 | 0 |
| AM+ | 0 | -10 | 10 |
| AM+ | -20 | -10 | 10 |
| NO-MAN | / | / | / |
| MIX | -20 | -10 | -10 |
| MIX | -20 | 0 | -10 |
| MIX | -20 | 10 | -10 |
| MIX | -20 | 10 | 0 |
| MIX | -20 | 10 | 10 |
| MIX | 0 | -10 | -10 |
| MIX | 0 | 10 | 10 |
| MIX | 20 | -10 | -10 |
| MIX | 20 | -10 | 0 |
| MIX | 20 | -10 | 10 |
| MIX | 20 | 0 | 10 |
| MIX | 20 | 10 | 10 |

**Table S2.**
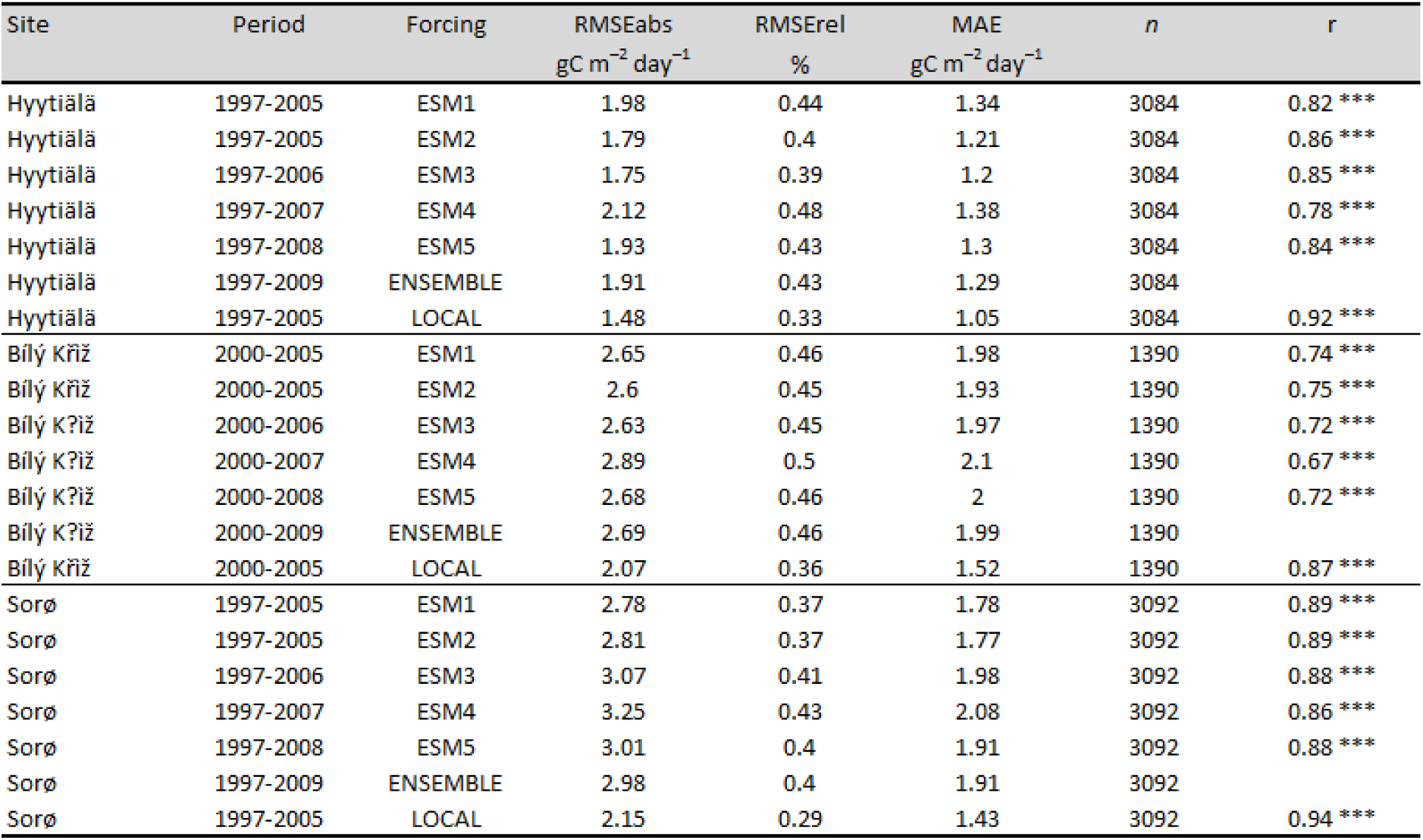
Performance statistics (Pearson correlation coefficient r, absolute and relative root mean square error RMSEabs (g C m^−2^ day^−1^) and RMSErel (%); mean absolute error, MAE (g C m^−2^ day^−1^) computed from daily series of model gross primary productivity, GPP against eddy covariance data when model is forced with measured (LOCAL) and modelled climate data (including the ensemble mean across ESMs climate data).

**Table S3.** Modeled and Estimated (bold) and modeled values of net primary productivity (NPP), plant autotrophic respiration (R_a_), and carbon use efficiency (CUE, i.e. NPP/GPP). Model output reports the annual average of the variables over the observation period. ‘LOCAL-Climate’ and ‘ESM-Climate’ refer to model output when driven by locally observed or climate-modeled data. (a = Wu et al., 2013; Knohl et al., 2008; Vanninen and Mäkelä 2005; Tang et al., 2014; b = Granier et al., 2008; Luyssaert et al., 2007; c = Xiao et al., 2003; Vanninen and Mäkelä 2005; Luyssaert et al., 2007; Tang et al., 2014)

|  | Sorø | Bílý Kříž | Hyytiälä |
| --- | --- | --- | --- |
| **NPP<sup>a</sup>** |  |  |  |
| **gC m<sup>-2</sup> year<sup>-1</sup>** | 708±65 | 611±45 | 434±55 |
| LOCAL | 860.35 | 547.31 | 454 |
| ESMs' ENSEMBLE | 734±32 | 517±28 | 416±26 |
| **R<sub>a</sub><sup>b</sup>** |  |  |  |
| **gC m<sup>-2</sup> year<sup>-1</sup>** | 751 ± 52 | 726±110 | 558±24.5 |
| LOCAL | 945.11 | 662.3 | 464.8 |
| ESMs' ENSEMBLE | 973±16 | 697±30 | 471±14 |
| **CUE<sup>c</sup>** |  |  |  |
| **adim.** | 0.45-0.50 | 0.45-0.5 | 0.32/0.45-0.54 |
| LOCAL | 0.47 | 0.45 | 0.49 |
| ESMs' ENSEMBLE | 0.42±0.01 | 0.42±0.01 | 0.47±0.01 |

**Table S4.** NPP and pCWS at the sites of Sorø, Bílý Křìž and Hyytiälä computed as percentage differences between the alternative management scenarios and the *Business-As-Usual* one are reported in parenthesis. Values refer to data averaged over the simulation period 2006-2099, across real and virtual standard climate for each RCPs. AM+ = more intensive than the *businnes-as-usual*; AM– = less intensive than the *businnes-as-usual*; NO- MAN = no management; RCPs = Representative Concentration Pathways. In bold the maximum and the minimum values across the RCPs and management type scenarios divided per site.

| Site | Management type | RCP | NPP<br>% | pCWS<br>% |
| --- | --- | --- | --- | --- |
| Sorø | AM+ | 2.6 | **-30.9** | **-6.7** |
|  | AM+ | 4.5 | -30.2 | -7.2 |
|  | AM+ | 6.0 | -30.3 | -7.2 |
|  | AM+ | 8.5 | -28.8 | -7.4 |
|  | AM- | 2.6 | **-2.2** | **-7.2** |
|  | AM- | 4.5 | -2.7 | -7.4 |
|  | AM- | 6.0 | -2.6 | -7.3 |
|  | AM- | 8.5 | -3.4 | -7.5 |
|  | NO-MAN | 2.6 | -18.8 | -30.0 |
|  | NO-MAN | 4.5 | -19.7 | -29.9 |
|  | NO-MAN | 6.0 | -19.3 | -29.9 |
|  | NO-MAN | 8.5 | -21.7 | -30.2 |
| Bílý Kříž | AM+ | 2.6 | -22.6 | 1.5 |
|  | AM+ | 4.5 | -21.6 | 1.9 |
|  | AM+ | 6.0 | -21.7 | 1.4 |
|  | AM+ | 8.5 | -19.1 | **3.0** |
|  | AM- | 2.6 | **-8.2** | -8.9 |
|  | AM- | 4.5 | -10.3 | -10.0 |
|  | AM- | 6.0 | -9.5 | -9.6 |
|  | AM- | 8.5 | -11.2 | -10.4 |
|  | NO-MAN | 2.6 | -36.6 | -35.8 |
|  | NO-MAN | 4.5 | -39.6 | -38.1 |
|  | NO-MAN | 6.0 | -39.0 | -38.0 |
|  | NO-MAN | 8.5 | **-41.7** | **-40.0** |
| Hyytiälä | AM+ | 2.6 | -37.7 | -7.0 |
|  | AM+ | 4.5 | -37.6 | -7.4 |
|  | AM+ | 6.0 | **-37.9** | -7.7 |
|  | AM+ | 8.5 | -37.1 | -7.1 |
|  | AM- | 2.6 | 9.2 | -3.8 |
|  | AM- | 4.5 | 7.7 | -4.1 |
|  | AM- | 6.0 | 7.6 | **-3.9** |
|  | AM- | 8.5 | 5.6 | -4.2 |
|  | NO-MAN | 2.6 | **21.6** | -21.6 |
|  | NO-MAN | 4.5 | 16.4 | -22.5 |
|  | NO-MAN | 6.0 | 15.9 | -22.1 |
|  | NO-MAN | 8.5 | 9.0 | **-23.7** |

**Table S5.**
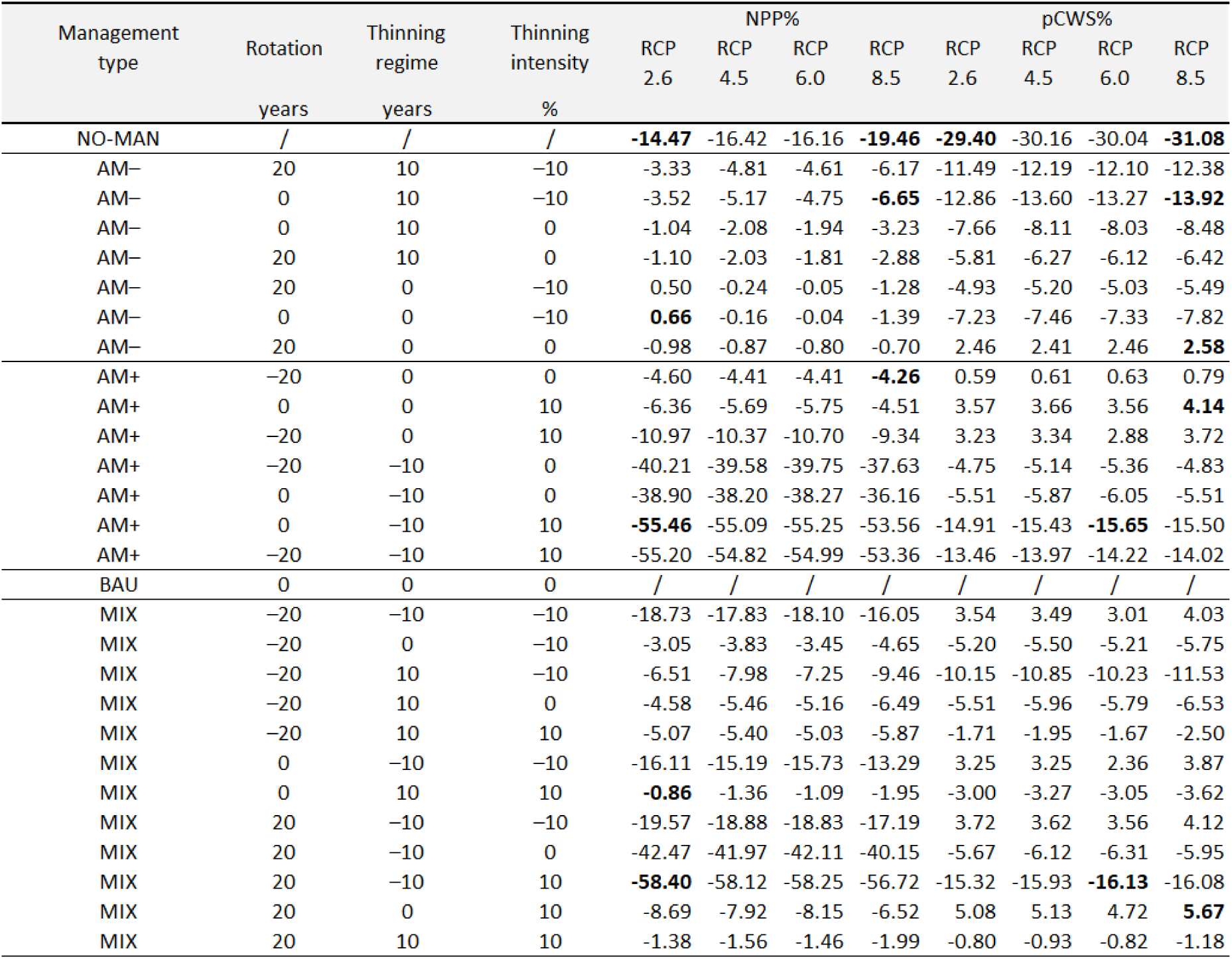
Percentage of changes for Mean NPP (net primary productivity) and pCWS (potential C woody stocks) over the period 2006-2099, between each alternative management scenario, the NO-MAN (no-management) scenario and the *business-as-usual scenarios* (BAU). Values refer to data averaged across real and virtual stands, species and climate for each RCPs. Positive values indicate a positive effect of one management scheme compared to BAU. ESMs = Earth System Models; RCPs = Representative Concentration Pathways, *r* = rotation turn, *th* = thinning frequency, *thi*= thinning intensity expressed as year or % differences compared to the BAU scheme, MIX = mixed scenario. In bold the maximum and the minimum values across the RCPs and management type scenarios.

**Table S6.** Factorial analysis (values range from 0 to 100) for the described variables (GPP, R_a_, NPP, CUE, NPP_woody_ and potential Carbon woody stock) across sites and the mean between sites.

| Factor | Bílý Kříž - GPP | Bílý Kříž - NPP | Bílý Kříž - $R_s$ | Bílý Kříž - CUE | Bílý Kříž - NPP<sub>woody</sub> | Bílý Kříž - pCWS |
| --- | --- | --- | --- | --- | --- | --- |
| ESM | 11.79 | 19.33 | 7.75 | 11.93 | 13.63 | 11.90 |
| ESM-RCP | 3.78 | 12.47 | 3.76 | 11.57 | 8.39 | 1.16 |
| ESM-RCP-Stand | 2.94 | 2.37 | 2.51 | 2.28 | 2.13 | 1.75 |
| ESM-Stand | 2.00 | 2.00 | 1.84 | 1.40 | 1.98 | 1.11 |
| RCP | 1.12 | 3.40 | 1.34 | 4.57 | 2.19 | 0.25 |
| RCP-Stand | 0.80 | 0.76 | 0.71 | 0.60 | 0.67 | 0.48 |
| Stand | 77.56 | 59.67 | 82.10 | 67.64 | 71.01 | 83.35 |
| Error | 0.00 | 0.00 | 0.00 | 0.00 | 0.00 | 0.00 |
| | Hyytiälä - GPP | Hyytiälä - NPP | Hyytiälä - $R_s$ | Hyytiälä - CUE | Hyytiälä - NPP<sub>woody</sub> | Hyytiälä - pCWS |
| ESM | 3.21 | 5.15 | 4.31 | 9.55 | 7.04 | 2.67 |
| ESM-RCP | 0.90 | 4.45 | 1.16 | 7.08 | 6.46 | 0.23 |
| ESM-RCP-Stand | 0.15 | 0.53 | 0.18 | 0.38 | 0.91 | 0.05 |
| ESM-Stand | 0.39 | 0.92 | 0.51 | 0.57 | 1.37 | 0.26 |
| RCP | 3.11 | 2.41 | 3.65 | 4.11 | 3.17 | 0.36 |
| RCP-Stand | 0.33 | 0.51 | 0.44 | 0.19 | 0.72 | 0.05 |
| Stand | 91.90 | 86.04 | 89.76 | 78.12 | 80.33 | 96.38 |
| Error | 0.00 | 0.00 | 0.00 | 0.00 | 0.00 | 0.00 |
| | Sorø - GPP | Sorø - NPP | Sorø - $R_s$ | Sorø - CUE | Sorø - NPP<sub>woody</sub> | Sorø - pCWS |
| ESM | 21.07 | 21.54 | 15.11 | 8.47 | 14.13 | 15.03 |
| ESM-RCP | 10.07 | 17.88 | 5.22 | 9.42 | 11.84 | 0.42 |
| ESM-RCP-Stand | 0.31 | 0.68 | 0.22 | 0.49 | 0.60 | 0.03 |
| ESM-Stand | 0.83 | 1.33 | 0.94 | 0.85 | 1.45 | 0.27 |
| RCP | 8.69 | 8.90 | 6.99 | 3.25 | 6.45 | 1.54 |
| RCP-Stand | 0.41 | 0.64 | 0.51 | 0.37 | 0.76 | 0.07 |
| Stand | 58.64 | 49.03 | 71.01 | 77.15 | 64.77 | 82.65 |
| Error | 0.00 | 0.00 | 0.00 | 0.00 | 0.00 | 0.00 |
| | ALL - GPP | ALL - NPP | ALL - $R_s$ | ALL - CUE | ALL - NPP<sub>woody</sub> | ALL - pCWS |
| ESM | 12.02 | 15.34 | 9.06 | 9.98 | 11.60 | 9.87 |
| ESM-RCP | 4.92 | 11.60 | 3.38 | 9.36 | 8.90 | 0.60 |
| ESM-RCP-Stand | 1.13 | 1.19 | 0.97 | 1.05 | 1.22 | 0.61 |
| ESM-Stand | 1.07 | 1.42 | 1.09 | 0.94 | 1.60 | 0.54 |
| RCP | 4.31 | 4.90 | 4.00 | 3.98 | 3.94 | 0.72 |
| RCP-Stand | 0.51 | 0.63 | 0.55 | 0.39 | 0.72 | 0.20 |
| Stand | 76.03 | 64.91 | 80.96 | 74.31 | 72.04 | 87.46 |
| Error | 0.00 | 0.00 | 0.00 | 0.00 | 0.00 | 0.00 |

